# GSK3β phosphorylation catalyzes the aggregation of Tau into Alzheimer’s disease-like amyloid strain

**DOI:** 10.1101/2023.12.19.572292

**Authors:** Pijush Chakraborty, Alain Ibáñez de Opakua, Jeffrey A. Purslow, Simon A. Fromm, Debdeep Chatterjee, Milan Zachrdla, Sambhavi Puri, Benjamin Wolozin, Markus Zweckstetter

## Abstract

The pathological deposition of proteins is a hallmark of several devastating neurodegenerative diseases. These pathological deposits comprise aggregates of proteins that adopt distinct structures named strains. However, the molecular factors responsible for the formation of distinct aggregate strains are unknown. Here we show that the serine/threonine kinase GSK3β catalyzes the aggregation of the protein tau into an Alzheimer’s disease-like amyloid strain. We demonstrate that phosphorylation by GSK3β, but not by several other kinases, promotes the aggregation of full-length tau through enhanced phase separation into gel-like condensate structures. Cryo-electron microscopy further reveals that the amyloid fibrils formed by GSK3β-phosphorylated tau adopt a fold comparable to that of paired helical filaments isolated from the brains of AD patients. Our results elucidate the intricate relationship between post-translational modification and the formation of tau strains in neurodegenerative diseases.

## Introduction

More than 30 neurodegenerative diseases are characterized by the pathological aggregation of the microtubule-associated protein tau into insoluble deposits^1,2^. In Alzheimer’s disease (AD), tau deposition correlates with disease progression^3^. Tau deposits in AD are composed of amyloid fibrils^4,5^. Tau is hyperphosphorylated in the diseased brain of AD and other tauopathies^3,6^. Phosphorylation has thus been suggested to modulate the formation of pathological tau aggregates and subsequent neuronal dysfunction^7^. Additionally, post-translational modifications, potentially together with so-far unknown co-factors, were suggested to drive the tau protein into distinct amyloid fibril structures^8,9^. These distinct tau fibril structures are named strains and are associated with different tauopathies^9–11^. However, the molecular factors responsible for the formation of specific tau strains remain unknown.

Phosphorylation is the most abundant modification observed in tau that regulates the physiological activities of tau by modulating its binding to microtubules and other cellular compartments^12^. Many serine/threonine kinases phosphorylate tau in the brain, including glycogen synthase kinase 3 (GSK3), cyclin-dependent kinase 5 (CDK5), microtubule affinity-regulating kinases (MARK), calmodulin-dependent protein kinase II (CaMKII), extracellular signal-regulated kinase (ERK) and cyclic AMP-dependent protein kinase (PKA)^12,13^. Furthermore, tyrosine kinases, such as FYN, SYK, and ABL, can phosphorylate tau’s tyrosine residues^12^. Phosphorylation of tau inside the microtubule-binding domain reduces its affinity towards the negatively charged microtubules^14,15^. Under pathological conditions, approximately 40-50 out of 85 potential phosphorylation sites in tau have been identified to undergo phosphorylation^16^.

A growing number of studies suggest that biomolecular condensation may contribute to the pathogenesis of neurodegenerative diseases by acting as an intermediate or transition state between the monomeric protein and its pathological aggregated state^17,18^. Consistent with this hypothesis, condensate formation can enhance the aggregation of intrinsically disordered proteins into amyloid fibrils^19,20^. Post-translational modifications including phosphorylation modulate biomolecular condensation either by promoting the phase separation of intrinsically disordered proteins or by inhibiting it^21^. For example, phosphorylation in the proline-rich domain of tau promotes the protein’s phase separation, while acetylation blocks tau phase separation^22–24^. However, little is known about the relationship between phosphorylation, tau condensation and aggregation, and their impact on the formation of distinct tau fibril structures in neurodegenerative diseases.

## Results

### Kinase-specific phosphorylation patterns of tau

To identify the tau phosphorylation patterns of different kinases, we performed individual phosphorylation reactions of the full-length 2N4R isoform of tau (441 residues; further referred to as tau) *in vitro*. For the phosphorylation reactions, we selected the six serine/threonine kinases GSK3β, CDK5, ERK2, MARK2cat, PKA and CaMKII, as well as the tyrosine kinase C-Abl (Fig. 1a). To assess primed phosphorylation, we also phosphorylated tau with two or three kinases in a step-wise manner: CDK5 followed by GSK3β, PKA followed by GSK3β, CDK5 followed by GSK3β and MARK2cat, as well as PKA followed by GSK3β and subsequently MARK2cat (Fig. 1a). To determine the phosphorylation patterns of tau, the phosphorylated samples were loaded in an SDS-PAGE gel (Supplementary Fig. 1). Subsequently, the tau bands observed in the SDS-PAGE gel were analyzed by mass spectrometry. The extent of phosphorylation for each residue was determined by the ratio between the number of detected peptides containing a particular phosphorylated residue and the total number of peptides detected containing the same residue.

**Fig. 1.**
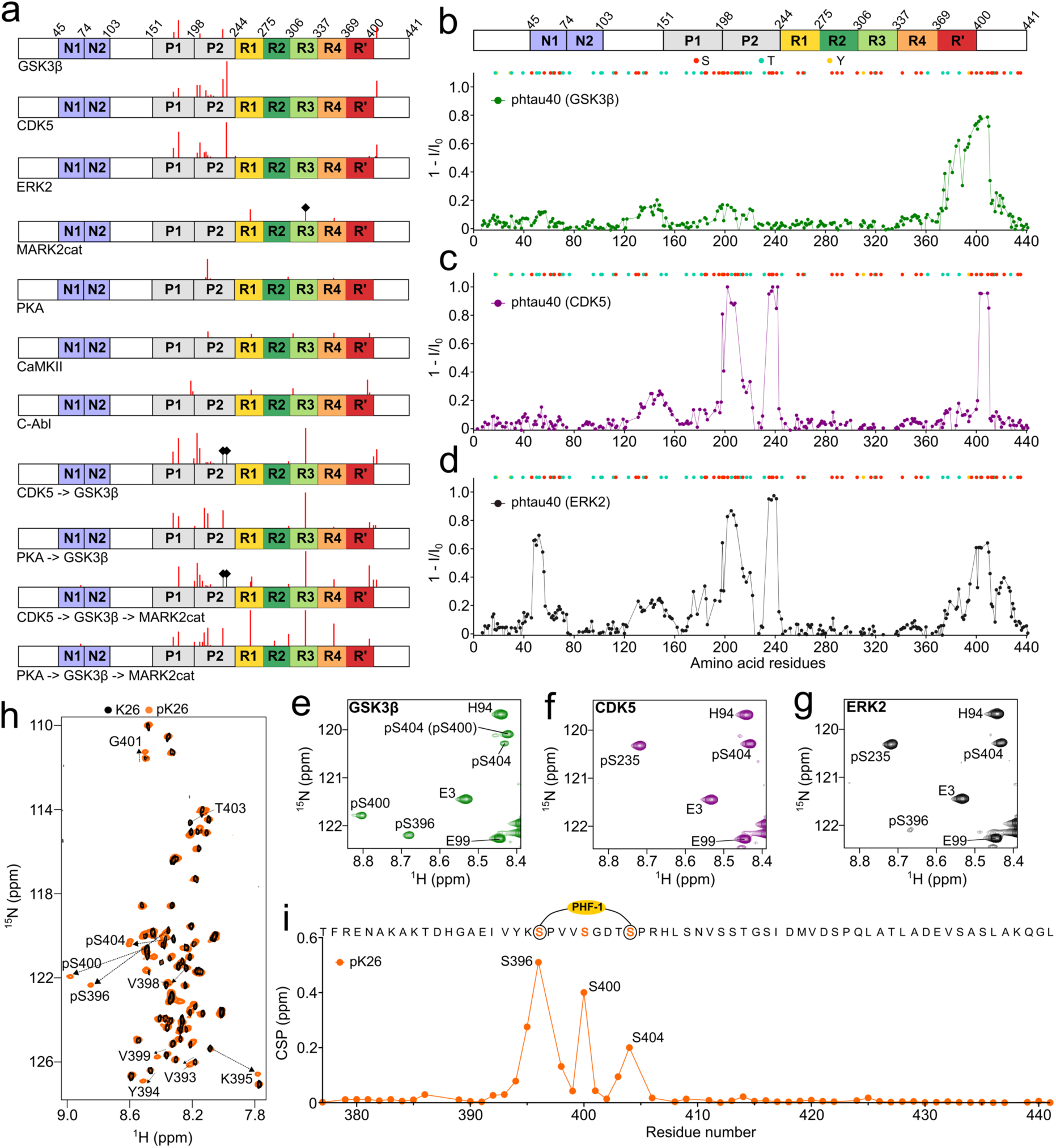
Kinase-specific phosphorylation patterns of tau. **a,** Domain diagram of full-length tau. The residues phosphorylated by different kinases and detected by mass spectrometry are indicated with red bars. The lengths of the red ticks are adjusted based on the percentage of phosphorylation calculated by dividing the number of peptides detected containing a particular residue by the total number of peptides detected containing the same residue. The black diamond-headed bars refer to previously reported phosphorylation sites that were not detected due to the absence of peptides in the mass spectrometry experiment. **b-d**, Residue-specific intensity changes observed in 2D ^1^H-^15^N HSQC spectra of tau upon phosphorylation by GSK3β (b), CDK5 (c) and ERK2 (d). I and I_0_ are the cross-peak intensities of the phosphorylated and unmodified tau, respectively. The location of serine, threonine and tyrosine residues is indicated above. **e-g**, Signals of the cross-peaks of phosphorylated serine residues in the ^1^H-^15^N HSQC spectra of phosphorylated tau appeared due to the phosphorylation by GSK3b (e), CDK5 (f), and ERK2 (g). **h**, ^1^H-^15^N HSQC spectra of either unmodified (black) or GSK3β-phosphorylated (orange) C-terminal fragment of tau comprising residues 369 to 441 (referred to as K26). The cross-peaks of the phosphorylated residues and the residues shifted due to nearby phosphorylation are shown in the spectrum. **i**, Residue-specific chemical-shift perturbation (CSP) observed in the ^1^H-^15^N HSQC spectra of K26 upon phosphorylation by GSK3β. The PHF-1 epitope is marked in the amino acid sequence displayed above.

The analysis provided insights into the specific tau phosphorylation patterns of the seven kinases. In the case of GSK3β, we detected phosphorylation of S404, as well as phosphorylation of some residues in the proline-rich domain (T175, T181, T205, T231) (Fig. 1a). However, detection of a very small number of peptides from tau’s C-terminus precluded confirmation of phosphorylation of S396 and S400, the two other serine residues in the C-terminus of tau that were previously shown to be also phosphorylated by GSK3β^25^. In the case of CDK5, we detected phosphorylation of the C-terminal site S404, as well as phosphorylation of T175, T181, S202, T205, T212, S214, T217, T220, T231and S235 in the proline-rich domain (Fig. 1a). The kinase ERK2 phosphorylated S396, T403 and S404 in the C-terminus of tau, as well as other sites, including T175, T181, S202, T205, S210, T212, S214, T217, S235 and T245.

The other three tested kinases (PKA, MARK2cat and CaMKII) also phosphorylated residues in the aggregation-prone repeat-domain of tau (Fig. 1a), but the sites differed from the specific sites phosphorylated by GSK3β, CDK5 and ERK2. For example, a phosphorylation reaction of 15 minutes in the presence of PKA results in the phosphorylation of S305 and S356 in repeat 2 and 4, respectively, together with phosphorylation of S214. Similarly, the catalytic domain of the kinase MARK2 (MARK2cat) phosphorylated S262 and S356. The phosphorylation at S324 by MARK2cat^26^ was not detected because of the absence of peptides from that region. In agreement with previous reports^25^, CaMKII phosphorylated S214, T263, and S356 of tau. A small number of peptides with phosphorylated Y310 and Y394 were also detected after CaMKII phosphorylation (Fig. 1a). The tyrosine kinase C-Abl phosphorylated Y197, Y310, and Y394 (phosphorylation of Y18 and Y29 was not observed due to the detection of few peptides from the N-terminus of tau). Sequential phosphorylation of tau by two or three kinases lead to an abundant phosphorylation in both the proline-rich domain and the repeat domain, with most peptides detected for phosphorylated S324 (Fig. 1a).

### GSK3β most efficiently phosphorylates tau’s PHF-1 epitope

To gain further insights into the sites and degree of phosphorylation catalyzed by the serine/threonine kinases GSK3β, CDK5 and ERK2, we used NMR spectroscopy. We phosphorylated ^15^N-labeled 2N4R tau in the presence of either kinase and recorded two-dimensional (2D) ^1^H-^15^N HSQC NMR spectra of the phosphorylated proteins. Comparison of the cross-peaks intensities in the phosphorylated samples to the unmodified sample revealed that the three kinases efficiently phosphorylated residues in the C-terminal domain of tau (Fig. 1b-d). The detection of newly appearing cross-peaks of the phosphorylated residues confirmed that GSK3β efficiently phosphorylates S396 and S404, i.e. the two residues that comprise the PHF-1 epitope^27^, as well as S400 (Fig. 1e). The extent of phosphorylation of S404 (∼75 %) and S400 (∼57 %) by GSK3β was calculated by comparing the intensity of the phosphorylated cross-peak to the unmodified cross-peak. On the other hand, we were unable to calculate the extent of phosphorylation of S396 due to signal overlap of unmodified S396 with T231. Notably, the NMR-based intensity analysis also shows that the other sites (T175, T181, T205, T231) in the proline-rich domain of tau, for which phosphorylation was detected by mass spectrometry (Fig. 1a), are only very weakly phosphorylated by GSK3β. As these sites are weakly phosphorylated by GSK3β, we also did not detect signals of the phosphorylated peaks of these residues. GSK3β thus most efficiently phosphorylates the PHF-1 epitope in the C-terminus of tau.

We further confirmed phosphorylation of S396, S400 and S404 by GSK3β-catalyzed phosphorylation of a C-terminal fragment of tau comprising residues 369 to 441 (referred to as K26) and subsequent analysis by NMR. The analysis revealed that GSK3β indeed phosphorylates S396, S400 and S404 in the C-terminal domain (Fig. 1h,i; Supplementary Fig. 2), where the extent of phosphorylation were determined to be 56 %, 59 %, and 81 % respectively.

The NMR analysis also showed that ERK2 phosphorylated both S404 and S396 but not S400 (Fig. 1g). Notably, the cross-peak of phosphorylated S396 was weaker than in case of GSK3β phosphorylation (Fig. 1g), suggesting that ERK2 is less efficient in phosphorylating S396 when compared to GSK3β. In the case of CDK5 phosphorylation, we did not detect a cross-peak of phosphorylated S396, indicating that CDK5 only phosphorylates S404 in the C-terminal domain of tau (Fig. 1f).

Besides phosphorylation at the C-terminus, CDK5 and ERK2 (GSK3β to a lesser degree) effectively phosphorylated residues in the proline-rich domain of tau (Fig. 1b-d), broadly in agreement with the results from mass spectrometry (Fig. 1a). The ERK2 kinase also phosphorylated some residues (most likely T50^28^) in the N-terminal regions between residue 40 and 80 (Fig. 1d). The phosphorylation of these residues were not detected by mass spectrometry, because of the detection of very few peptides from the N-terminus of tau. The combined data show that GSK3β and ERK2, but not the other tested kinases, phosphorylate the tau epitope (pS396 & pS404) that is recognized by the monoclonal antibody PHF-1 ^27^.

### GSK3β phosphorylation accelerates tau amyloid formation

We next asked whether the distinct phosphorylation patterns imprinted by different kinases differentially modulate the aggregation of tau into insoluble deposits. To gain insight into this question, we aggregated the unmodified and phosphorylated tau proteins using our previously developed co-factor-free *in vitro* aggregation assay^29^. We discovered distinct phosphorylation-specific changes in tau amyloid formation: only GSK3β-phosphorylated tau aggregated faster than the unmodified protein (Fig. 2a,b; Supplementary Fig. 3). Other tau proteins phosphorylated by a single kinase displayed slower aggregation when compared to unmodified tau. ERK2-phosphorylated tau was most similar to unmodified tau in its aggregation kinetics, followed by PKA-, CDK5- and CamKII-phosphorylated tau (Fig. 2a,b; Supplementary Fig. 3). C-Abl-phosphorylated tau displayed even slower aggregation kinetics, but tau aggregation was most impaired when phosphorylated by the catalytic domain of MARK2. Interestingly, stepwise phosphorylation by two kinases delayed tau aggregation when compared to phosphorylation by the respective individual kinases. Phosphorylation by two kinases and additionally by MARK2 largely blocked tau fibrillization during the time of incubation (Fig. 2a,b; Supplementary Fig. 3). Thus, GSK3β phosphorylation selectively accelerates tau amyloid formation.

**Fig. 2.**
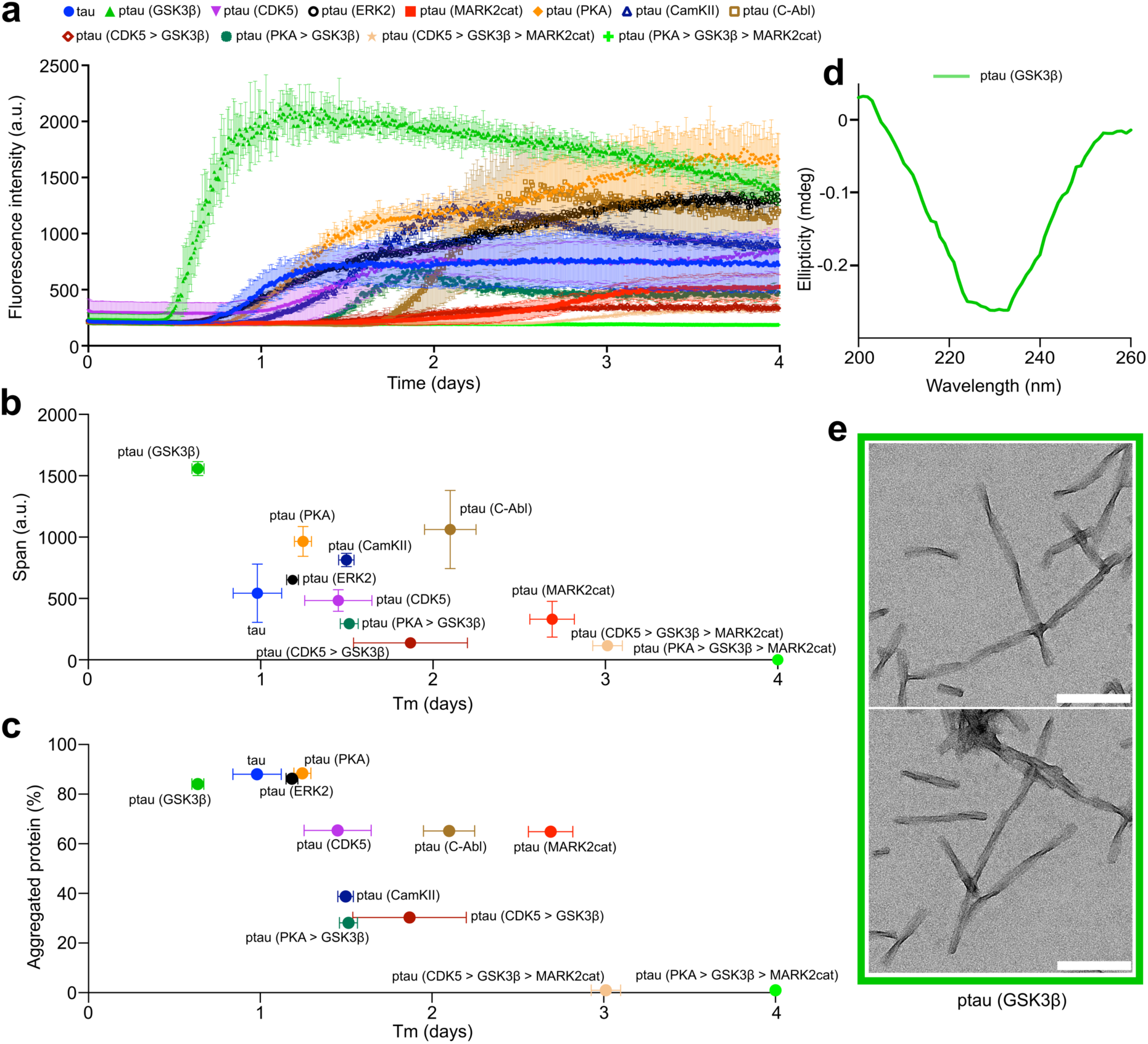
Acceleration of tau aggregation by GSK3β phosphorylation. **a**, Aggregation kinetics of 25 µM unmodified tau and tau phosphorylated by different kinases. Error bars represent the std of three independently aggregated samples. **b**, ThT-intensity span vs. half time of aggregation (Tm) of unmodified and phosphorylated tau proteins. Error bars represent the std of three independently aggregated samples. **c**, Amount of protein aggregated vs. half time of aggregation (Tm) of unmodified and phosphorylated tau proteins. The amount of aggregated protein was calculated by comparing the intensity of the supernatant (SN) band (after pelleting down the fibrils) to the tau monomer band as shown in Supplementary Fig. 4. Error bars represent the std of the half-time of aggregation of three independently aggregated samples. **d**, CD spectrum of fibrils formed by GSK3β-phosphorylated tau at the end of the aggregation assay. **e**, Negative-stain EM of fibrils formed by GSK3β-phosphorylated tau. Scale bars, 200 nm.

In addition to phosphorylation-induced changes in the aggregation kinetics, we observed different final fluorescence intensities of the amyloid-specific dye thioflavin-T (ThT) for tau proteins phosphorylated by different kinases (Fig. 2a,b; Supplementary Fig. 3). Most of the phosphorylated tau proteins displayed higher ThT intensity at the end of aggregation when compared to the unmodified protein, apart from tau proteins that were phosphorylated by multiple kinases (Fig. 2a,b).

To evaluate whether the higher ThT intensity is caused by an increased amount of aggregated protein, we centrifuged the aggregated samples at the end of the incubation period. Subsequently, the amount of residual protein was analyzed by loading the supernatants into an SDS-PAGE gel (Supplementary Fig. 4). Comparison of the band intensity of the supernatants with the unmodified monomeric protein revealed that a similar amount of protein was aggregated for unmodified tau and tau phosphorylated by the kinases GSK3β, ERK2 and PKA (Fig. 2c). Approximately 80-90% of the monomeric proteins were aggregated into insoluble deposits (Fig. 2c). In contrast, phosphorylation of tau by CDK5, C-Abl, and MARK2cat decreased the amount of aggregated protein to ∼60% (Fig. 2c). The amount of aggregated protein further decreased when tau was phosphorylated by CamKII or in the case of sequential phosphorylation with two kinases. Finally, the sequential phosphorylation of tau in the presence of three kinases, either PKA or CDK5 followed by GSK3β and MARK2cat, abolished tau fibrillization. Thus, the higher ThT intensity observed for most of the fibrils formed by phosphorylated tau proteins is likely caused by differences in ThT binding, indicating the formation of distinct amyloid fibril structures by phosphorylated tau proteins when compared to unmodified tau.

To gain insight into the impact of GSK3β-phosphorylation - the only phosphorylation that accelerated tau amyloid formation when compared to the unmodified protein (Fig. 2a-c) - on tau amyloid structure, we recorded circular dichroism (CD) and negative-stain electron microscopy (EM) images (Fig. 2d,e). The CD spectrum confirmed the presence of β-structure, which is characteristic of disease-associated amyloid fibrils (Fig. 2d). In the EM images, we observed separated fibrils with a pronounced twist (Fig. 2e). These twisted fibrils differ from the fibrils formed by unmodified tau which were mostly straight (Supplementary Fig. 5).

### Phosphorylation at PHF-1 epitope catalyzes the condensation of tau

To understand whether the accelerated aggregation of GSK3β-phosphorylated tau is linked to the formation of condensates, we imaged the GSK3β-phosphorylated tau samples by differential interference contrast (DIC) microscopy prior to starting the aggregation assay. Under this condition, i.e., after incubation for ten minutes at room temperature in the aggregation assay buffer, we observed the formation of condensates in the GSK3β-phosphorylated tau sample (Fig. 3b). Fluorescence microscopy confirmed that these condensates are indeed formed by GSK3β-phosphorylated tau (Fig. 3b). In contrast, the unmodified tau did not form visible condensates in these conditions (Fig. 3g).

**Fig. 3.**
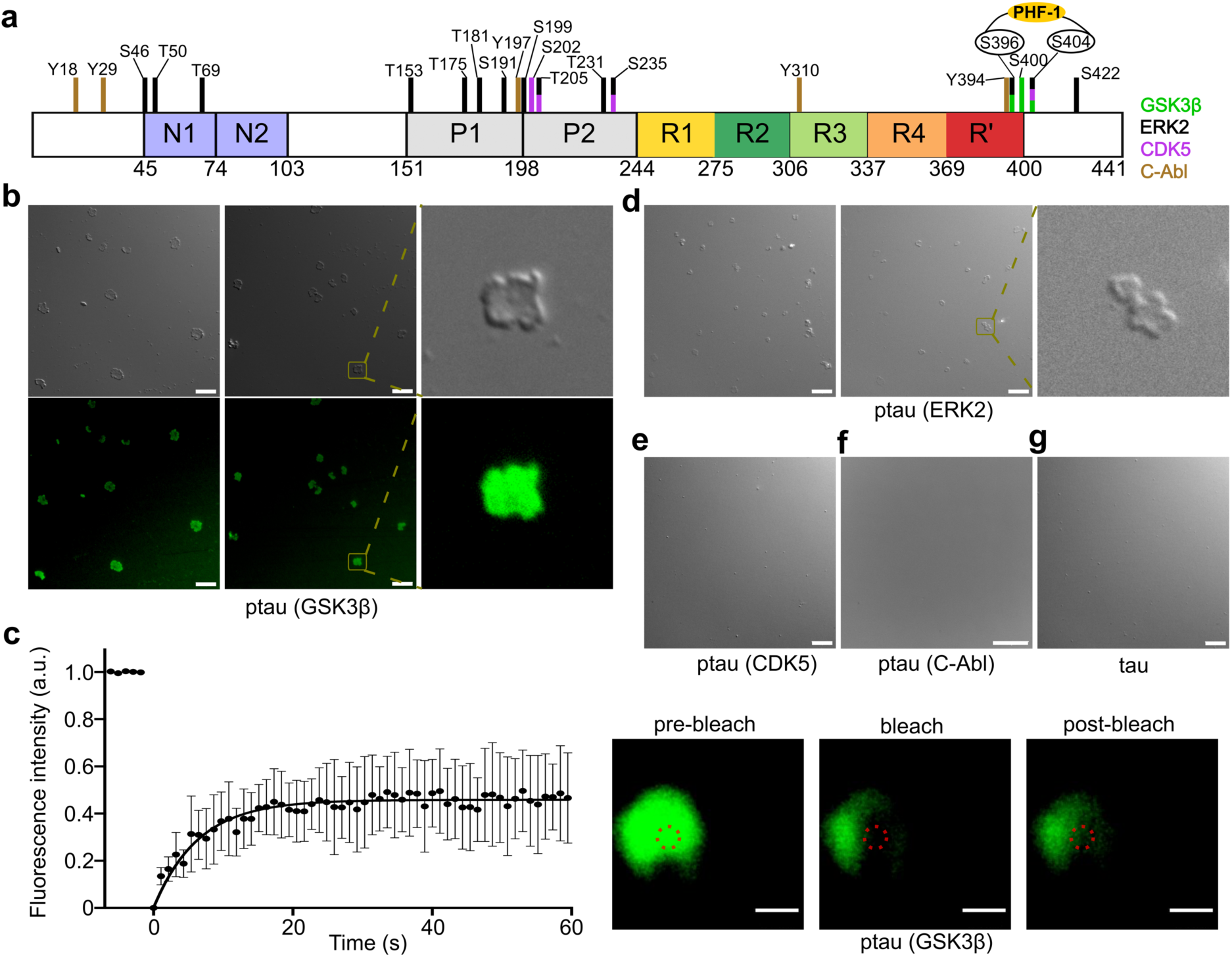
GSK3β phosphorylation promotes tau condensation. **a**, Domain diagram of 2N4R tau. Residues that undergo phosphorylation in the presence of GSK3β, ERK2, CDK5, and C-Abl are marked with green, black, purple, and dark-yellow colored bars, respectively. S396 and S404, which are phosphorylated by GSK3β, form the epitope that is recognized by the phosphorylation-specific antibody PHF-1. **b**, DIC and fluorescence microscopy of the condensates formed by 25 µM GSK3β-phosphorylated tau at room temperature in the aggregation assay buffer (25 mM HEPES, 10 mM KCl, 5 mM MgCl_2_, 3 mM TCEP, 0.01 % NaN_3_, pH 7.2). A zoomed-in view of the condensate is shown to the right. Micrographs are representative of three independent biological replicates. Scale bar, 10 µm **c**, Fluorescence recovery after photobleaching (FRAP) experiment of the condensates formed by GSK3β-phosphorylated tau shown in (b). Error bars represent std of averaged three curves for each time point. Representative micrographs of the condensate before bleaching, after bleaching, and at the end of recovery are displayed to the right. Scale bar, 1 µm. **d**, DIC microscopy of the condensates formed by 25 µM ERK2-phosphorylated tau at room temperature in the aggregation assay buffer. A zoomed-in view of the condensate is shown to the right. Micrographs are representative of three independent biological replicates. Scale bar, 10 µm **e,f,g**, DIC microscopy of 25 µM CDK5-phosphorylated tau (e), C-Abl-phosphorylated tau (f), and unmodified tau (g) at room temperature in the aggregation assay buffer. Micrographs are representative of three independent biological replicates. Scale bar, 10 µm.

Closer inspection of the DIC images showed that the condensates formed by GSK3β-phosphorylated tau did not possess the typical spherical shape of liquid droplets, but displayed a more “squeezed” appearance. These condensates were also stained by the amyloid-binding dye Thioflavin-T (Supplementary Fig. 6). To gain insight into the diffusive properties of these condensates, we used fluorescence recovery after photobleaching. Although part of the fluorescence recovered rapidly in a few seconds, an immobile fraction of 50-80 % remained (Fig. 3c). This suggests that the material properties inside the GSK3β-phosphorylated tau condensates differ from liquid droplets and contain both more diffusive and gel-like tau protein.

The major GSK3β-induced phosphorylation sites in tau are located at the C-terminus (S396, S400, and S404) (Fig. 1b,e, Fig. 3a). Similar to GSK3β, ERK2 (S396, S404), and CDK5 (S404) also phosphorylate tau at the C-terminus (Fig. 1c,d,f,g; Fig. 3a). To elucidate the relationship between phosphorylation site and condensate formation, we imaged ERK2-phosphorylated as well as CDK5-phosphorylated tau with DIC microscopy prior to starting the aggregation assay (Fig. 3d,e). In the aggregation assay buffer, we observed that ERK2-phosphorylated tau forms condensates similar to that of GSK3β-phosphorylated tau, but not CDK5-phosphorylated tau (Fig. 3d,e). As an additional control, we asked whether other types of phosphorylation may promote condensate formation. However, tau phosphorylated by the tyrosine kinase C-Abl, which phosphorylates multiple of the tyrosine residues of tau including Y394 at the C-terminus, did not form condensates under the same conditions (Fig. 3f). This suggests that phosphorylation at only S404 is not sufficient, but it requires phosphorylation of both S396 and S404, that is the PHF-1 epitope, to promote tau condensation.

### Tau droplets mature into gel-like structures upon GSK3β-phosphorylation

Crowding agents promote the liquid-liquid phase separation (LLPS) of unmodified tau^22,30,31^. In agreement with these reports, we observed the formation of spherical, highly mobile droplets of unmodified tau in the presence of 10 % dextran (Fig. 4a). We then added fluorescently labeled GSK3β to the pre-formed tau droplets and observed that the kinase rapidly enters the droplets (Fig. 4a). GSK3β thus concentrates inside the tau droplets where it might phosphorylate tau.

**Fig. 4.**
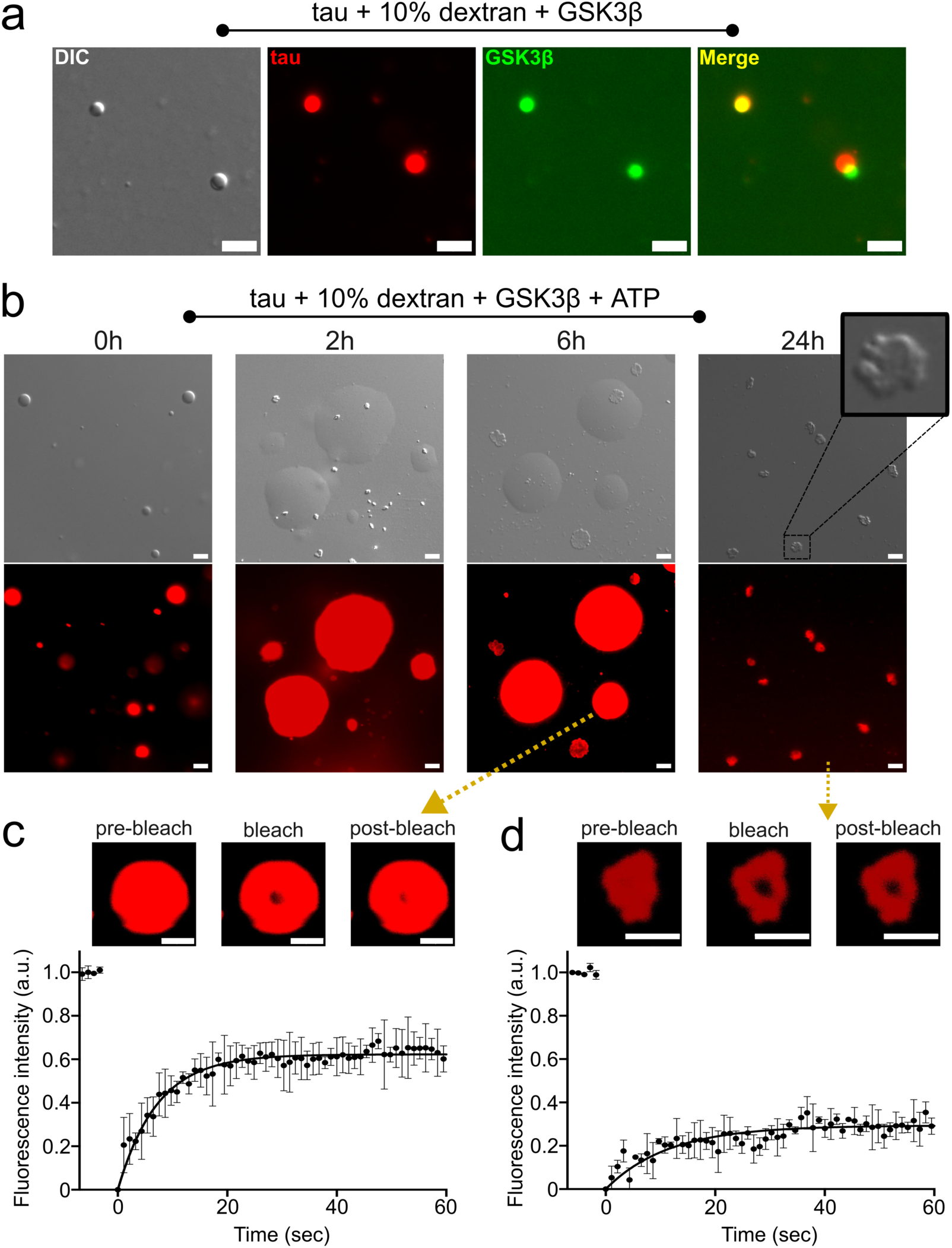
Maturation of tau droplets into gel-like structures upon GSK3β phosphorylation. **a**, DIC and fluorescence microscopy of tau droplets induced by the addition of 10 % dextran at room temperature in 25 mM HEPES, 5 mM MgCl_2_, pH 7.2 buffer. Fluorescently labeled GSK3β partitioned into the droplets. The partially merged image is due to the mobility of the droplets, i.e., the droplets moved from one place to another place by the time the images were taken using the red and green channel of the microscope. Micrographs are representative of three independent biological replicates. Scale bar, 5 µm. **b**, DIC and fluorescence microscopy of the tau droplets induced by the addition of 10 % dextran at room temperature in 25 mM HEPES, 5 mM MgCl_2_, pH 7.2 buffer in the presence of 0.02 mg/ml unlabeled GSK3β, and 1 mM ATP. The sample was incubated for 24 hours. A zoomed-in view of the condensate formed after 24 hours is shown. Micrographs are representative of three independent biological replicates. Scale bar, 5 µm. **c**, FRAP experiment of the larger fused condensates of tau in the presence of GSK3β and ATP after incubation for six hours. Error bars represent std of averaged three curves for each time point. Representative micrographs of the condensate before bleaching, after bleaching, and at the end of recovery are displayed on the top. Scale bar, 5 µm. **d**, FRAP experiment of the gel-like condensates of tau in the presence of GSK3β and ATP after incubation for one day. Error bars represent std of averaged three curves for each time point. Representative micrographs of the condensate before bleaching, after bleaching, and at the end of recovery are displayed on the top. Scale bar, 5 µm.

To assess whether GSK3β phosphorylates tau and changes the morphology of tau droplets, we repeated the experiment but now added unlabeled GSK3β as well as ATP to the pre-formed tau droplets. During the first two hours of incubation, we observed mobile tau droplets that fused and formed larger droplets (Fig. 4b). After six hours of incubation, we again observed the larger mobile tau droplets, but additionally some smaller “squeezed”-like condensates appeared (Fig. 4b). After one day of incubation, we observed the presence of only such gel-like condensates in the solution (Fig. 4b). They were morphologically similar to the condensates formed by the GSK3β-phosphorylated tau in absence of a crowding agent (Fig. 3b). In contrast, we did not observe such gel-like condensates upon the incubation of tau droplets only in the presence of GSK3β, i.e. under non-phosphorylating conditions without ATP (Supplementary Fig. 7).

Mass spectrometry confirmed that the tau sample was indeed phosphorylated under the condition of LLPS (Supplementary Fig. 8). Notably, the total degree of phosphorylation was higher when phosphorylation occurred under the conditions of LLPS when compared to GSK3β-induced phosphorylation in the dispersed phase (Supplementary Fig. 8), in agreement with the acceleration of phosphorylation in condensed states of tau^32^.

To get further insight into the dynamics inside the tau condensates, we performed FRAP experiments. Photobleaching the larger tau condensates formed with GSK3β but without ATP, i.e. in non-phosphorylating conditions, after incubation for six hours resulted in an immobile fraction of ∼20% (Supplementary Fig. 7b). A prolonged incubation of one day resulted in an immobile fraction between 30% to 40% (Supplementary Fig. 7c). In contrast, photobleaching the larger tau condensates formed in the presence of both GSK3β and ATP after incubation for six hours revealed the presence of immobile fractions of ∼40 % (Fig. 4c). Additionally, photobleaching the gel-like condensates, which formed only in the presence of both GSK3β and ATP after an incubation of one day, resulted in an immobile fraction of ∼80 % (Fig. 4d). The ∼80% immobile fraction is at the upper end of immobile fractions observed in the condensates formed by GSK3β-phosphorylated tau in the aggregation assay buffer without crowding agents (Fig. 3c). This may result from a longer incubation time, different buffer conditions, or the presence of dextran in the current experiments. Taken together, the data show that tau droplets mature into gel-like structures upon GSK3β-phosphorylation.

### GSK3β-phosphorylated tau folds into AD-like amyloid

The above experiments show that the specific phosphorylation of the PHF-1 epitope by GSK3β promotes the condensation of tau into gel-like structures that further convert into amyloid fibrils under conditions of agitation. To characterize the stable core region of the fibrils formed by GSK3β-phosphorylated tau, we combined pronase digestion with mass spectrometry. We first digested GSK3β-phosphorylated tau fibrils by pronase, centrifuged the sample, then loaded the pellet on an SDS-PAGE (Supplementary Fig. 9a). Subsequently, the tau band at ∼12 kDa was cut and digested by trypsin, followed by detection of the peptides using an ESI mass spectrometer. Analysis of the number of detected peptides as a function of sequence revealed peptides belonging to residues ∼280 to ∼400 (Supplementary Fig. 9b). The pronase-resistant core of GSK3β-phosphorylated tau fibrils thus comprises residues ∼280 to ∼400, which is comparable to the core of tau fibrils extracted by sarkosyl from the brain tissues of AD patients^33^.

To understand the structure of GSK3β-phosphorylated tau fibrils at higher resolution, we utilized cryo-EM. The cryo-EM images of the GSK3β-phosphorylated tau revealed the presence of both straight and twisted filaments (Fig. 5a). The average crossover distance of the twisted fibrils was ∼1550 Å (Fig. 5b). 3D reconstructions of the twisted filaments were calculated to an overall resolution of ∼5 Å using helical reconstruction in RELION^34,35^ (Supplementary Fig. 10 and Supplementary Table 1).

**Fig. 5.**
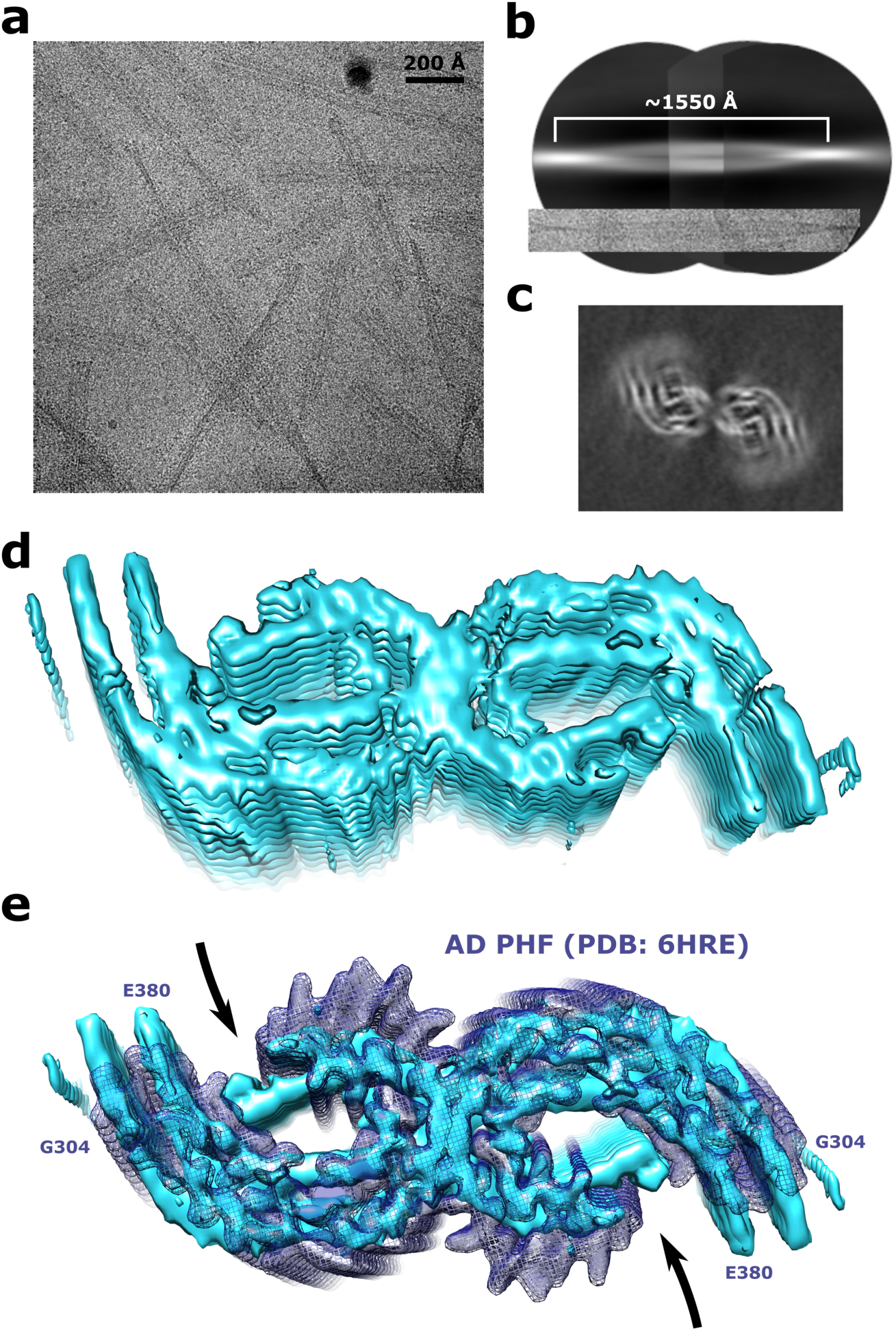
Cryo-EM of GSK3β-phosphorylated tau fibrils. **a**, Cryo-electron micrograph of GSK3β-phosphorylated tau fibrils. **b,** Reconstruction of a GSK3β phosphorylated tau fibril from low-resolution and big box 2D classes for crossover estimation. **c,** Cross-section of the cryo-EM map of the GSK3β-phosphorylated tau fibril after 3D refinement. **d,** Cryo-EM density map of GSK3β phosphorylated tau fibrils. **e,** Cryo-EM density map of GSK3β-phosphorylated tau fibrils (cyan surface) compared with the paired helical filament from sporadic Alzheimer’s disease brain (dark blue isomesh; PDB id: 6HRE).

The GSK3β-phosphorylated tau fibrils are composed of two protofilaments with closed C-shaped subunits (Fig. 5c,d). The protofilaments are related by C2 symmetry and the successive rungs of β-sheets in each protofilament are related through helical symmetry with a rise of 4.75 Å and twist of approximately −0.55° (Supplementary Fig. 10a and Supplementary Table 1).

The cryo-EM analysis shows that the structure adopted by the GSK3β-phosphorylated tau fibrils is similar to that of paired helical filaments (PHF) derived from the brain of AD patients (Fig. 5e). Indeed, the location of the rigid core is comparable, the individual filaments fold into a C-shaped conformation, and the two filaments have a similar relative arrangement (Fig. 5e). While the current resolution does not allow further analysis, the protofilaments of the *in vitro* aggregated GSK3β-phosphorylated tau fibrils may adopt a more closed C-shaped conformation when compared to the AD PHFs (Fig. 5e), which are related by a pseudo 2-1 screw symmetry^33,36^. The GSK3β-phosphorylated tau fibrils may thus be more similar to the PHFs extracted from the extracellular vesicles isolated from the brains of people with Alzheimer’s disease (Supplementary Fig. 11).

## Discussion

The proline-directed serine/threonine kinase GSK3β is closely associated with the pathological phosphorylation of tau in AD^37–39^. In AD patients, GSK3β co-localizes with neurofibrillary tangles^40,41^ and concentrates in the frontal cortex of AD brains^38^. In animal models, increased GSK3β activity leads to tau hyperphosphorylation and downstream neurodegeneration^37^. Accordingly, an isoform-selective decrease of GSK3β reduces synaptic tau phosphorylation, transcellular spreading, and aggregation^39^. Here we showed that GSK3β most efficiently phosphorylates the PHF-1 epitope of tau, catalyzes the phase separation of tau into gel-like condensates, and selectively accelerates *in vitro* aggregation of tau into amyloid fibrils that adopt a fold similar to that found in PHFs extracted from AD patient brains.

Using our recently described co-factor-free aggregation assay of full-length tau, we showed that GSK3β but none of the other five tested kinases, nor the combinations of two or three kinases, promote the aggregation of tau into amyloid fibrils (Fig. 2). Using a combination of mass spectrometry and NMR spectroscopy, we linked the accelerated aggregation of GSK3β-phosphorylated tau to the ability of GSK3β to efficiently phosphorylate S396 and S404 of tau, which are recognized by the monoclonal antibody PHF-I (Fig. 1). While the kinase ERK2 also phosphorylates both S396 and S404, the degree of S396 phosphorylation was lower when compared to GSK3β-phosphorylation. The efficient phosphorylation of S396 and the additional phosphorylation of S400, which is only phosphorylated by GSK3β, may contribute to the GSK3β-specific acceleration of tau aggregation. The three serine residues at the C-terminus (S396, S400, S404) that are phosphorylated by GSK3β, are also abnormally hyperphosphorylated in the brain of Alzheimer’s disease patients^42^. Notably, phosphorylation at the N-terminus and in particular in the repeat domain delayed tau aggregation (Fig. 1). Our study also shows that GSK3β without priming by another kinase only weakly phosphorylates residues T181 and T231, sites in the proline-rich domain that are likely phosphorylated early in the disease process. This indicates that it may be important to further investigate priming of GSK3β phosphorylation by a variety of kinases and its impact on tau condensation and aggregation.

Our data further reveals that under the conditions of aggregation, tau phosphorylated by GSK3β (without priming by other kinases) forms condensates with gel-like properties. In contrast, unmodified tau did not form condensates under the same conditions (Fig. 3). The condition of condensate formation may increase the kinetics of aggregation because the high tau concentration inside the condensates may lead to faster aggregation on the surface of the condensate^19,20^. Additionally, we found that phosphorylation by GSK3β is stronger in conditions of phase separation when compared to the dispersed phase. Phosphorylation by GSK3β and tau condensation may thus reinforce each other. The connection between tau condensation and accelerated aggregation was further supported by our observation that other types of phosphorylated tau protein exhibit delayed aggregation when compared to unmodified tau and do not form condensates (Fig. 3e,f,g). Phosphorylation of tau by ERK2 elicited condensate formation without enhanced aggregation, which contrasts with observations for the other kinase/tau pairings (Fig. 2, 3). These data point to additional mechanisms – in particular to the regulatory role of phosphorylation in the proline-rich domain – that differentially regulate tau condensation and aggregation.

In brain-derived extracts from AD patients, filaments comprising only two C-shaped protofilaments of tau have so far been observed^33,36^ (Fig. 5e). Additionally, a truncated fragment of 4R tau comprising residues 297 to 391 was recently shown to aggregate *in vitro* into a fibril structure comprising two protofilaments that is highly similar to that of *ex vivo* PHFs^43^. The same study also reported that another C-terminal pseudo-phosphorylated 4R tau fragment (S396D, S400D, T403D, and S404D) comprising residues from 297 to 441 adopts a single protofilament with the AD fold^43,44^. Although this study reported the *in vitro* aggregation of a 4R tau fragment into the AD fold comprising two protofilaments, the tau species that are deposited in the brains of AD patients are predominantly the full-length isoforms of both 4R and 3R tau^7,43,45^. Here, we showed that the full-length 4R isoform of tau phosphorylated at the C-terminus by the AD-associated kinase GSK3β aggregates *in vitro* into filaments that are composed of two C-shaped protofilaments with a fold that is similar to that of *ex vivo* PHFs (Fig. 5). However, the C-shaped protofilaments formed by GSK3β-phosphorylated 4R tau *in vitro* adopt a more closed/compact shape as compared to the AD PHFs (Fig 5e). In addition, the resolution of the fibril structure of GSK3β-phosphorylated tau is currently not sufficient to perform a more detailed comparison. Notably, a recent study reported that the tau PHF derived from the extracellular vesicles of an AD patient’s brain adopts a more compact fold as compared to the AD PHF isolated from total brain homogenates (Supplementary Fig. 11)^46^. The compactness of the AD PHF isolated from the extracellular vesicles is due to the presence of an additional co-factor between two positively-charged residues (R349 and K375). Consequently, the presence of different co-factors in different regions of the brain may drive tau to adopt an even more compact protofilament fold in certain subtypes of AD. Further work is also needed to understand the contribution of 3R tau isoforms to the formation of the AD tau strain.

Taken together, our study identifies specific C-terminal phosphorylation of tau as a major molecular factor leading to the formation of AD-like tau amyloid structure. It further strengthens the critical role played by the serine/threonine kinase GSK3β in the pathogenesis of AD.

## Methods

### Protein purification

To prepare unlabeled 2N4R tau protein, a single colony from the LB-agar plate was taken and grown overnight in 50 mL LB medium supplemented with 100 µg/mL ampicillin at 37 °C. 22 mL of the overnight culture were transferred to 1L LB medium supplemented with 100 µg/mL ampicillin and allowed to grow until an OD_600_ of 0.8-0.9 was reached. Subsequently, the cells were induced with 0.5 mM IPTG and expressed for 1 hour.

To obtain uniformly ^15^N-labeled 2N4R tau, cells were grown in 8 L LB until an OD_600_ of 0.6-0.8 was reached, then centrifuged at low speed (5,000 g), washed with 1X M9 salts, and resuspended in 2 L M9 minimal medium supplemented with 1 g/L ^15^NH_4_Cl as the only nitrogen source. After 1 hour, the cells were induced with 0.5 mM IPTG and expressed overnight at 37 ^0^C.

After harvesting, cell pellets were resuspended in lysis buffer (20 mM MES (pH 6.8), 1 mM EGTA, 2 mM DTT) complemented with protease inhibitor mixture, 0.2 mM MgCl_2_, lysozyme, and DNAse I. Subsequently, cells were disrupted with a French pressure cell press (in ice-cold conditions to avoid protein degradation). NaCl was added to a final concentration of 500 mM, and lysates were boiled for 20 minutes. Denatured proteins were removed by ultracentrifugation with 127,000 g at 4 °C for 30 minutes. To precipitate the DNA, 20 mg/mL streptomycin sulfate was added to the supernatant and incubated for 15 minutes at 4 °C followed by centrifugation at 15,000 g for 30 minutes. The pellet was discarded, and tau protein was precipitated by adding 0.361 g/mL ammonium sulfate to the supernatant, followed by centrifugation at 15,000 g for 30 minutes. The pellet containing tau protein was resuspended in buffer A (20 mM MES (pH 6.8), 1 mM EDTA, 2 mM DTT, 0.1 mM PMSF, 50 mM NaCl) and dialyzed against the same buffer (buffer A) to remove excess salt. The next day, the sample was filtered and applied to an equilibrated ion-exchange chromatography column (Mono S 10/100 GL, GE Healthcare), and weakly bound proteins were washed out with buffer A. Tau protein was eluted with a linear gradient of 60 % final concentration of buffer B (20 mM MES pH 6.8, 1 M NaCl, 1 mM EDTA, 2 mM DTT, 0.1 mM PMSF). Protein samples were concentrated by ultrafiltration (5 kDa Vivaspin, Sartorius) and further purified by reverse phase chromatography using a preparative C4 column (Vydac 214 TP, 5 µm, 8 x 250 mm) in an HPLC system coupled with ESI mass spectrometer. Protein purity was confirmed using mass spectrometry, and the purified protein was lyophilized and re-dissolved in the buffer of interest. To prepare unlabeled GSK3β (kinase domain), a construct of human GSK3β (residues 35-386) with an N-terminal His-tag was cloned into a pET28a vector. A thrombin consensus sequence (LVPR/GS, where / signifies the cleaved peptide bond) was positioned between the His-tag and GSK3β components of the construct, leaving a four-residue overhang following the removal of the His-tag. The cloned vector was transformed into *E. coli* strain BL21-AI and subsequently plated onto an LB-agar plate treated with 50 µg/mL of Kanamycin for selection. A seed culture was produced by growing a single colony in 100 mL of Luria Broth (LB) medium overnight at 37 °C and 230 rpm. 20 mL of the seed culture was then inoculated into 1L of LB medium and grown at 37 °C and 110 rpm to an OD_600_ ∼0.4, where the temperature was then reduced to 16 °C. At an OD_600_ ∼0.6, recombinant protein expression was induced by the addition of 2 g/L (0.2%) of arabinose and 0.4 mM of Isopropyl ß-D-1-thiogalactopyranoside (IPTG). After 18 h of induction at 16 °C and 110 rpm, cells were harvested by centrifugation for 30 min at 5,000 g. The supernatant was discarded, and the cell pellet resuspended in lysis buffer (50 mM HEPES (pH 7.2), 300 mM NaCl, 5% glycerol, 1 mM phenylmethylsulfonyl fluoride (PMSF), 20 µg/mL DNase, 0.1 mg/mL lysozyme, and EDTA-free protease inhibitor cocktail). The cell suspension was lysed on ice by sonication and subsequently centrifuged at 50,000 g for 30 min. The supernatant was decanted from cellular debris and passed through a 0.45 µM filter. The supernatant was loaded onto a 5 mL HisTrap HP prepacked column (Cytiva), washed with buffer A (50 mM HEPES (pH 7.2), 300 mM NaCl, 5% glycerol) and eluted using a gradient of buffer B (50 mM HEPES (pH 7.2), 300 mM NaCl, 5% glycerol, and 500 mM imidazole). Elution fractions were analyzed by SDS-polyacrylamide gel (SDS-PAGE) electrophoresis and all fractions containing target protein coalesced. The sample was then diluted with dilution buffer (50 mM HEPES (pH 7.2) and 5% glycerol) to reduce the salt concentration and loaded onto a Mono S 10/100 GL column (GE Healthcare). The column was washed with buffer C (50 mM HEPES (pH 7.2), 100 mM NaCl, and 5% glycerol), before eluting the target protein using a gradient of buffer D (50 mM HEPES (pH 7.2), 100 mM NaCl, 5% glycerol, and 1 M NaCl). The resulting elution was analyzed by SDS-PAGE electrophoresis, where the purist fractions were combined and concentrated using a 10K MWCO Vivaspin membrane filter (Sartorius). 72 units of thrombin per mg of protein were added and the sample was dialyzed overnight at 4 °C into 50 mM HEPES (pH 7.2), 150 mM NaCl, 5% glycerol, and 2 mM 2-Mercaptoethanol (BME). Following dialysis, the sample was loaded onto 1 mL of Ni-NTA beads equilibrated with buffer A to remove any uncleaved protein. The protein was then further purified using a HiLoad 16/600 Superdex 200 Size Exclusion Chromatography (SEC) column equilibrated with 20 mM MES (pH 6.5), 250 mM NaCl, 5% glycerol, and 2 mM BME. Following the complete purification, the target protein was calculated to be ∼94% pure by analyzing an SDS-PAGE gel loaded with varying concentrations of protein.

### Phosphorylation of tau

Phosphorylation reactions of tau in the presence of a single kinase were performed according to published protocols (MARK2cat – Schwalbe et al.^26^, GSK3β and CaMKII – Ukmar-Godec et al.^25^, PKA – Leroy et al.^47^, ERK2 – Qi et al.^28^, C-Abl – Savastano et al.^48^). The reaction time and the concentration of kinases were chosen such that the phosphorylation occurs predominantly on the sites that are phosphorylated most efficiently by the respective kinases (i.e., less phosphorylation on minor sites).

Phosphorylation of 200 µM tau was performed in the presence of 2.5 µM of MARK2cat WT kinase, 5 mM ATP, 1 mM Benzamidine, 2 mM EGTA, 1 mM PMSF in 25 mM Tris, 100 mM NaCl, 5 mM MgCl_2_, 1 mM DTT, pH 8.0 buffer for a duration of 12 hours at 30 °C with 300 rpm shaking in an Eppendorf thermomixer.

Phosphorylation of 200 µM tau was performed with either 0.02 mg/ml of GSK3β (kinase domain) or 0.02 mg/ml of CDK5 and in the presence of 2 mM ATP, 5 mM EGTA, 1 mM PMSF in 40 mM HEPES, 5 mM MgCl_2_, 2 mM DTT, pH 7.4 buffer for a duration of 12 hours at 30 °C with 300 rpm shaking in an Eppendorf thermomixer.

Phosphorylation of 200 µM tau was performed in the presence of 1 µM PKA, 5 mM ATP, 5 mM EGTA, 1 mM PMSF in 50 mM HEPES, 12.5 mM MgCl_2_, 50 mM NaCl, 5 mM DTT, pH 8.0 buffer for a duration of 15 minutes at 30 °C with 300 rpm shaking in an Eppendorf thermomixer.

Phosphorylation of 200 µM tau was performed in the presence of 1 µM ERK2, 2.5 mM ATP, 2 mM EGTA, 1 mM PMSF in 50 mM HEPES, 12.5 mM MgCl_2_, 50 mM NaCl, 2 mM DTT, pH 8.0 buffer for a duration of 3 hours at 37 °C with 300 rpm shaking in an Eppendorf thermomixer.

Phosphorylation of 200 µM tau was performed in the presence of 1 µM C-Abl, 5 mM ATP, 2 mM EGTA, 1 mM PMSF in 40 mM HEPES, 5 mM MgCl_2_, 2 mM DTT, pH 7.4 buffer for a duration of 12 hours at 30 °C with 300 rpm shaking in an Eppendorf thermomixer.

Phosphorylation of 200 µM tau was performed in the presence of 0.02 mg/ml CaMKII, 2 mM ATP, 1 mM PMSF, 1 mM CaCl_2_, 2 µM Calmodulin in 40 mM HEPES, 5 mM MgCl_2_, 2 mM DTT, pH 7.4 buffer for a duration of 12 hours at 30 °C with 300 rpm shaking in an Eppendorf thermomixer.

At the end of the phosphorylation reactions, the samples were boiled at 98 °C for 20 minutes to precipitate the kinases followed by centrifugation at 20,000 g in an Eppendorf centrifuge 5424. Next, the pellet was discarded, and the supernatant containing phosphorylated tau was dialyzed against the buffer of interest.

The phosphorylation reactions with multiple kinases were performed sequentially, i.e., the phosphorylation reaction was performed with one kinase followed by the next one. In all cases, the same protocol mentioned for each kinase was used.

### Aggregation assays

Unmodified as well as all phosphorylated tau samples were aggregated using the previously described co-factor free aggregation protocol^29^. Briefly, 25 µM of protein were aggregated at 37 °C in 25 mM HEPES, 10 mM KCl, 5 mM MgCl_2_, 3 mM TCEP, 0.01 % NaN_3_, pH 7.2 buffer (aggregation assay buffer) in a 96 well plate using a Tecan spark plate reader. Three PTFE beads along with double orbital shaking were used to promote fibrillization. Thioflavin-T (ThT) at a final concentration of 50 µM was used to monitor the aggregation kinetics.

### Circular dichroism

10 µL of 25 µM GSK3β-phosphorylated tau fibrils were pelleted down by centrifugation at 20,000 g using an Eppendorf centrifuge 5424. The supernatant was discarded, and the pellet was dissolved in 50 µL of distilled water. CD data were collected in a 0.02 cm pathlength cuvette using Chirascan-plus qCD spectrometer at 25 °C. Datasets were averaged from ten repeated measurements. The spectra were baseline corrected and smoothed with a window size of four.

130 µL of 5 µM unmodified or GSK3β-phosphorylated tau were incubated for 10 minutes at room temperature in the aggregation assay buffer. CD data were collected in a 0.1 cm pathlength cuvette using Chirascan-plus qCD spectrometer at 25 °C. Datasets were averaged from ten repeated measurements. The spectra were baseline corrected and smoothed with a window size of four.

### Protease digestion

50 µL of 0.8 mg/mL GSK3β-phosphorylated tau fibrils and 0.4 mg/ml of pronase (53702, Merck-Millipore), were incubated in the aggregation assay buffer for 30 minutes at 1,400 rpm shaking in an Eppendorf thermomixer at 37 °C. The pronase-resistant material was pelleted down by ultracentrifugation at 160,000 g for 30 minutes at 4 °C using a Beckman Coulter Optima MAX-UP ultracentrifuge. The supernatant was removed, and the pellet was redissolved in 10 µL of aggregation assay buffer and loaded in a 15 % SDS-PAGE gel. For mass spectrometry, the tau band from the SDS-PAGE gel was cut and digested by trypsin, followed by the detection of the peptides using an ESI mass spectrometer (Orbitrap Fusion Tribrid, Thermo Fischer Scientific).

### Microscopy

For the experiments reported in Fig. 3b-g, 25 µM of unmodified or GSK3β/ERK2/CDK5/C-Abl phosphorylated tau in the aggregation assay buffer (25 mM HEPES, 10 mM KCl, 5 mM MgCl_2_, 3 mM TCEP, 0.01 % NaN_3_, pH 7.2) were incubated at room temperature for ten minutes and 5 µL of the sample was loaded onto a glass side and covered with an 18 mm coverslip for DIC microscopy experiments. For fluorescence microscopy experiments, the GSK3β-phosphorylated tau was labeled with Alexa-fluor-488 C_5_ Maleimide (green) (Thermo Fischer Scientific) and 0.5 µL of the labeled protein were added to 15 µL of unlabeled protein, and from that mixture 5 µL of the sample were loaded onto a glass slide and covered with an 18 mm coverslip.

For the experiments reported in Fig. 4a, 25 µM tau were taken in 25 mM HEPES, 5 mM MgCl_2_, pH 7.2 buffer in the presence of 10 % dextran and incubated at room temperature for 5 minutes. Then, Alexa-fluor-488 (Thermo Fischer Scientific) labeled GSK3β (Kinase domain) was added to the solution at a final concentration of 0.02 mg/ml. 15 µL of the solution was then mixed with 0.5 µL of Alexa-fluor-594 (Thermo Fischer Scientific) labeled tau protein, and from that mixture 5 µL of the sample were loaded onto a glass slide and covered with an 18 mm coverslip.

For the experiments reported in Fig. 4b-d, 25 µM tau were taken in 25 mM HEPES, 5 mM MgCl_2_, pH 7.2 buffer in the presence of 10 % dextran and incubated at room temperature for 5 minutes. Then, unlabeled GSK3β (Kinase domain) and ATP were added to the solution at a final concentration of 0.02 mg/ml and 1 mM, respectively. 45 µL of the solution were then mixed with 1 µL of Alexa-fluor-594 (Thermo Fischer Scientific) labeled tau protein and incubated at room temperature for measurement by microscopy at different time points.

DIC and fluorescent micrographs were acquired on a Leica DM6B microscope with a 63x objective (water immersion) and processed using Fiji software (NIH).

### Fluorescence recovery after photobleaching (FRAP)

FRAP experiments were recorded using a Zeiss LSM880 confocal microscope using a 63x objective (oil immersion) and either a 488 or 561 argon laser line. A circular region was chosen in a region of homogenous fluorescence and bleached with up to twelve iterations of full laser power and then the recovery was imaged. Pictures were analyzed in FIJI software (NIH) and FRAP recovery curves were calculated using standard methods based on fluorescence intensities measured for pre-bleached/bleached, reference, and background ROI^49^. The pre-bleached/bleached ROI was a selected region in the droplet before/after bleaching; the reference ROI correspond to an area that did not experience bleaching, while the background ROI correspond to an area where no fluorescence was detected. The intensity of the background ROI was further subtracted from the pre-bleached/bleached and reference ROIs. Briefly, the FRAP recovery was calculated as:

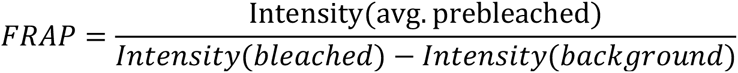

The value obtained was corrected by multiplying with the acquisition bleaching correction factor (ABCF) that was calculated according to

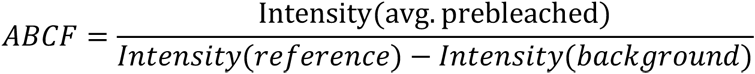

The curves were then normalized according to the following equation:

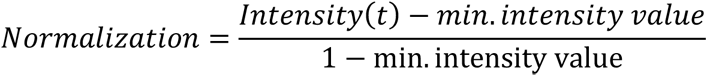

### NMR Spectroscopy

^1^H-^15^N HSQC spectra of uniformly ^15^N-labeled unmodified or GSK3β/CDK5/ERK2-phosphorylated tau (50 µM) were recorded in 50 mM NaP, 10 mM NaCl, 1 mM TCEP, pH 6.8 buffer at 278 K on an Avance III 900 MHz spectrometer (Bruker) using a 5 mm TCI (H/C/N) Cryoprobe. The spectra were collected with 40 scans per point (ns), and acquisition times td1 = 107.9 ms and td2 = 80.3 ms.

^1^H-^15^N HSQC spectra of uniformly ^13^C-^15^N-labeled unmodified K26 (30 µM) were recorded in 50 mM NaP, 10 mM NaCl, 1 mM TCEP, pH 6.8 buffer at 278 K on an Avance III 600 MHz spectrometer (Bruker) using a 5 mm QCI (H/C/N/F) Cryoprobe. The spectra were collected with 16 scans per point (ns), and acquisition times td1 = 43.8 ms and td2 = 155 ms.

^1^H-^15^N HSQC spectra of uniformly ^13^C-^15^N-labeled GSK3β-phosphorylated K26 (600 µM) were recorded in 50 mM NaP, 10 mM NaCl, 1 mM TCEP, pH 6.8 buffer at 278 K on an Avance III 600 MHz spectrometer (Bruker) using a 5 mm QCI (H/C/N/F) Cryoprobe. The spectra were collected with 8 scans per point (ns), and acquisition times td1 = 110.8 ms and td2 = 155 ms. The 3D HNCO spectra of the ^13^C-^15^N-labeled GSK3β-phosphorylated K26 was recorded on an Avance III 600 MHz spectrometer (Bruker) with 8 scans per point (ns), and acquisition times td1 = 47 ms, td2 = 22.5 ms, and td3 = 155.1 ms. A 3D HN(CA)CO spectrum was also recorded on the same spectrometer with 88 scans per point (ns), and acquisition times td1 = 23.5 ms, td2 = 20.7 ms, and td3 = 155.1 ms. A 3D HNCA spectrum of the ^13^C-^15^N-labeled GSK3β-phosphorylated K26 was recorded on an Avance Neo 800 MHz spectrometer (Bruker) using a 3 mm TCI (H/C/N) Cryoprobe with 16 scans per point (ns), and acquisition times td1 = 12.7 ms, td2 = 18.2 ms, and td3 = 118.8 ms. The 3D HN(CO)CA spectrum was recorded on the same spectrometer with 32 scans per point (ns), and acquisition times td1 = 12.7 ms, td2 = 12.9 ms, and td3 = 118.8 ms.

The chemical shift assignments of 2N4R tau had been previously reported^50^. The spectra were recorded using Topsin 3.6.2/4.0.3 software (Bruker) and analyzed with CCPNMR 2.4.2 software ^51^. The cross-peaks of the phosphorylated S396, S400, and S404 were assigned by performing sequential assignment of the GSK3β-phosphorylated K26 protein. These assignments of the phosphorylated residues were transferred to the phosphorylated full-length (2N4R) tau.

Residue-specific intensity ratios were calculated according to intensity ratio = 1− (I/I_0_), where I is the intensity of cross-peaks in the 2D ^1^H-^15^N HSQC spectrum of GSK3β/CDK5/ERK2-phosphorylated tau or GSK3β-phosphorylated K26 and I_0_ is the intensity of the cross-peaks of unmodified tau or K26.

The chemical shift perturbation (CSP) was calculated according to:

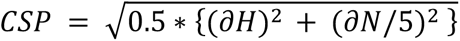

### In-gel digestion and extraction of peptides for mass spectrometry

To determine the phosphorylation patterns of tau, the phosphorylated samples were loaded in an SDS-PAGE gel (Supplementary Fig. 1). The respective bands from the SDS-PAGE gels were carefully cut and kept in an Eppendorf tube. To wash the gel pieces, 150 µL of water was added and incubated for 5 minutes at 26 °C with 1050 rpm shaking in a thermomixer. The gel pieces were spun down and the liquid was removed using thin tips (the same washing protocol was used in all subsequent steps with different solvents). The gel pieces were washed again with 150 µL acetonitrile. After washing, the gel pieces were dried for 5 minutes using a SpeedVacc vacuum centrifuge. To reduce disulfide bridges, 100 µL of 10 mM DTT was added to the gel pieces and incubated for 50 minutes at 56 °C followed by centrifugation and removal of liquid. The gel pieces were washed again with 150 µL of acetonitrile. To alkylate reduced cysteine residues, 100 µL of 55 mM iodoacetamide were added and incubated for 20 minutes at 26 °C with 1050 rpm shaking followed by centrifugation and removal of liquid. Subsequently, the gel pieces were washed with 150 µL of 100 mM NH_4_HCO_3_, and then twice with 150 µL of acetonitrile and dried for 10 minutes in a vacuum centrifuge. The gel pieces were rehydrated at 4 °C for 45 minutes by addition of small amounts (2-5 µL) of digestion buffer 1 (12.5 µg/mL trypsin, 42 mM NH_4_HCO_3_, 4 mM CaCl_2_). The samples were checked after every 15 minutes and more buffer was added in case the liquid was completely absorbed by the gel pieces. 20 µL of digestion buffer 2 (42 mM NH_4_HCO_3_, 4 mM CaCl_2_) were added to cover the gel pieces and incubated overnight at 37 °C.

To extract the peptides, 15 µl water was added to the digest and incubated for 15 minutes at 37 °C with 1050 rpm shaking followed by spinning down the gel pieces. 50 µl acetonitrile was added to the entire mixture and incubated for 15 minutes at 37 °C with 1050 rpm shaking. The gel pieces were spun down and the supernatant (SN1) containing the extracted peptides was collected. 30 µl of 5 % (v/v) formic acid was added to the gel pieces and incubated for 15 minutes at 37 °C with 1050 rpm shaking followed by spinning down. Again 50 µl acetonitrile were added to the entire mixture and incubated for 15 minutes at 37 °C with 1050 rpm shaking. The gel pieces were spun down and the supernatant (SN2) containing the extracted peptides was collected. Both supernatants (SN1 & SN2) containing the extracted peptides were pooled together and evaporated in the SpeedVacc vacuum centrifuge. The dried peptides were resuspended in 5 % acetonitrile and 0.1 % formic acid and analyzed using an Orbitrap Fusion Tribrid (Thermo Fischer Scientific) instrument. The MS data were analyzed using Scaffold 4 software.

### Negative-stain electron microscopy

40 µL of 25 µM GSK3β-phosphorylated tau fibrils were pelleted down by centrifugation at 20,000 g using an Eppendorf centrifuge 5424. The supernatant was discarded, and the pellet was redissolved in the aggregation assay buffer. Next, the fibrils were sonicated in a water bath (Bandelin sonorex) for 2 minutes. After sonication, 5.5 µL of fibril sample was mixed with 0.5 µL of 1 mg/mL pronase protease (53702, Merck-Millipore) followed by adsorbing onto carbon-coated copper grids. The samples were stained with 1% uranyl acetate solution and the images were taken with a Tietz F416 CMOS camera (TVIPS, Gauting, Germany) using a CM 120 transmission electron microscope (FEI, Eindhoven, The Netherlands).

### Cryo-electron microscopy

For cryo-EM, 5 µL of 25 µM GSK3β-phosphorylated tau fibrils were sonicated for 1 minute using a water bath sonicator (Bandelin Sonorex digitec) and then mixed with 1 µL of 1 mg/mL pronase (53702, Merck-Millipore). The mixture was then quickly added to the Quantifoil 2/1 grids and plunged-frozen after an incubation of 5 seconds using a Leica EM GP2 automatic plunge freezer. Cryo-electron microscopy data were acquired with a Titan Krios G4 transmission-electron microscope (Thermo Fisher) operated at 300 keV accelerating voltage. Images were recorded using a Falcon 4i direct electron detector with a calibrated pixel size of 0.934 Å on the specimen level. The slit width of the Selectris X energy filter (Thermo Fisher) was set to 10 eV and a 100 µm objective aperture was inserted. In total, 9,009 images with defocus values in the range of −0.9 µm to −1.9 µm were acquired in movie mode with 2.7 s acquisition time. The accumulated dose was approximately 40 electrons per Å^2^. The resulting dose-fractionated image stacks were subjected to beam-induced motion correction and CTF estimation.

Manual fibril picking was done with EMAN2 e2helixboxer^52^ to select an average of ∼5 segments per micrograph in 100 micrographs. The manual picking was used to train a model and pick the rest of the micrographs with crYOLO^53^ with an inter-box distance of 19 Å.

GSK3β phosphorylated tau fibrils were reconstructed using RELION-3.1.2^34,35^, following the helical reconstruction scheme. For an initial 2D classification, we extracted particle segments using a box size of 1536 pixels downscaled to 192 pixels. Best classes with a visible twist were selected and used to estimate a crossover of around 1550 Å (Fig. 5b), which is equivalent to a twist of around −0.55° for a 4.75 Å rise. For 3D classification, the segments after 2D classification were re-extracted without downscaling using a box size of 400 pixels. We performed several rounds of 3D classification starting from a 290 Å low-pass-filtered featureless cylinder and subsequent 3D refinements to optimize the helical parameters (rise of 4.77 Å and twist of −0.58°, reported in Supplementary Table 1) and applying C2 symmetry. Next, standard RELION post-processing with a soft-edged solvent mask that includes the central 20 % of the box height yielded the final post-processed map (sharpening B-factor of - 26.43 Å^2^). The resolution (3.85 Å) was estimated from the value of the FSC curve for two independently refined half-maps at 0.143. The estimated resolution is overestimated because of the high resolution in the Z axis and the approximated real resolution is around 5 Å (See Supplementary Fig. 10a,b).

## Acknowledgements

We thank the mass spectrometry facility of the Max Planck Institute for Multidisciplinary Sciences (MPI-NAT, Göttingen) for mass spectrometry and the EM facility of MPI-NAT for electron micrographs. We thank Kerstin Overkamp, MPI-NAT, for purifying tau constructs by reverse-phase chromatography. This work benefited from access to the EMBL Imaging Centre and has been supported by iNEXT-Discovery, project number 871037, funded by the Horizon 2020 program of the European Commission. We acknowledge the access and services provided by the Imaging Centre at the European Molecular Biology Laboratory (EMBL IC), generously supported by the Boehringer Ingelheim Foundation. M.Z. was supported by the European Research Council (ERC) under the EU Horizon 2020 research and innovation programme (grant agreement No. 787679).

## Author contributions

P.C. performed protein purification, phosphorylation reactions, aggregation assays, NMR experiments, DIC and fluorescence microscopy, protease digestion experiment, biophysical analysis, Cryo-EM sample preparation; A.I.O. solved the structure of GSK3β-ptau fibril by Cryo-EM; J.A.P. purified GSK3β (kinase domain) protein used to label with Alexa-Fluor-488; S.A.F. recorded the Cryo-EM dataset; D.C recorded the FRAP datasets; M.Zach. performed initial experiments on tau LLPS and GSK3β co-localization; S.P. performed experiments; B.W. supervised experiments; M.Z. supervised the project; P.C. and M.Z. designed the project and wrote the paper.

## Competing interests

The authors declare no competing financial interests.

## Notes

### Competing Interest Statement

The authors have declared no competing interest.

