## Supplementary information for "GSK3β phosphorylation catalyzes the aggregation of Tau into Alzheimer’s disease-like amyloid strain"

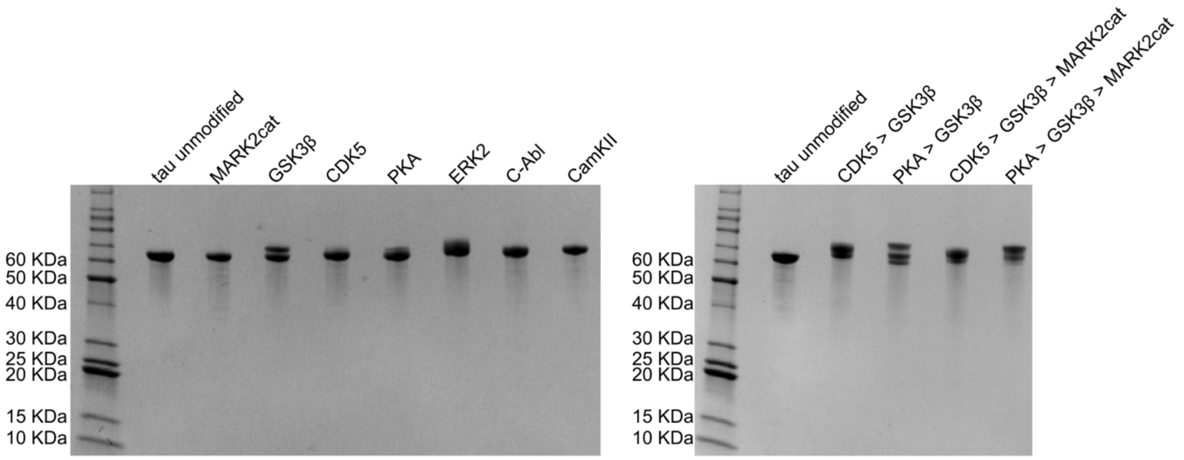


**Supplementary Fig. 1 | SDS-PAGE gel of unmodified and phosphorylated tau samples.** The shift in the tau bands toward higher molecular weight indicates the phosphorylation of the sample. These bands are cut and digested by trypsin and the resulting peptides are analyzed by mass spectrometry. The phosphorylated residues identified by mass spectrometry are shown in Fig. 1a.

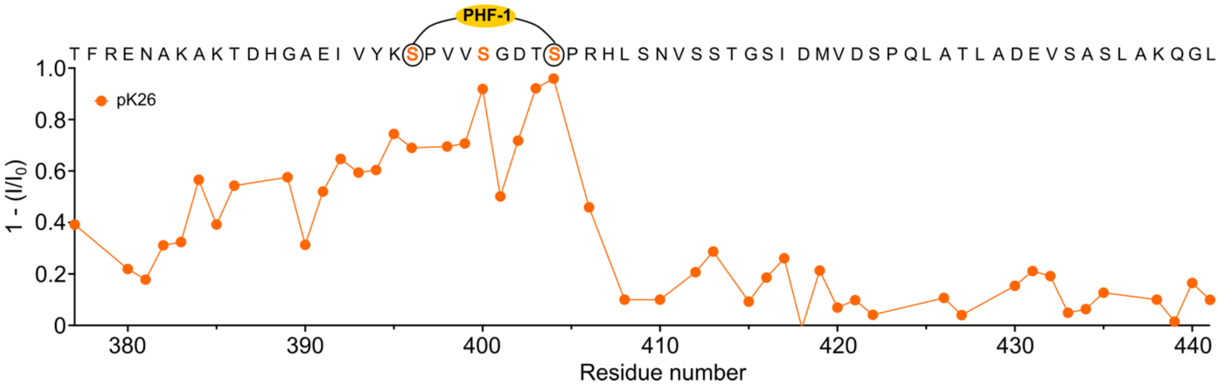


**Supplementary Fig. 2 | NMR spectroscopy of GSK3β-phosphorylated K26.** Residue-specific intensity changes observed in the $^{1}H$-$^{15}N$ HSQC spectra (Fig. 1h) of K26 upon phosphorylation by GSK3β kinase.

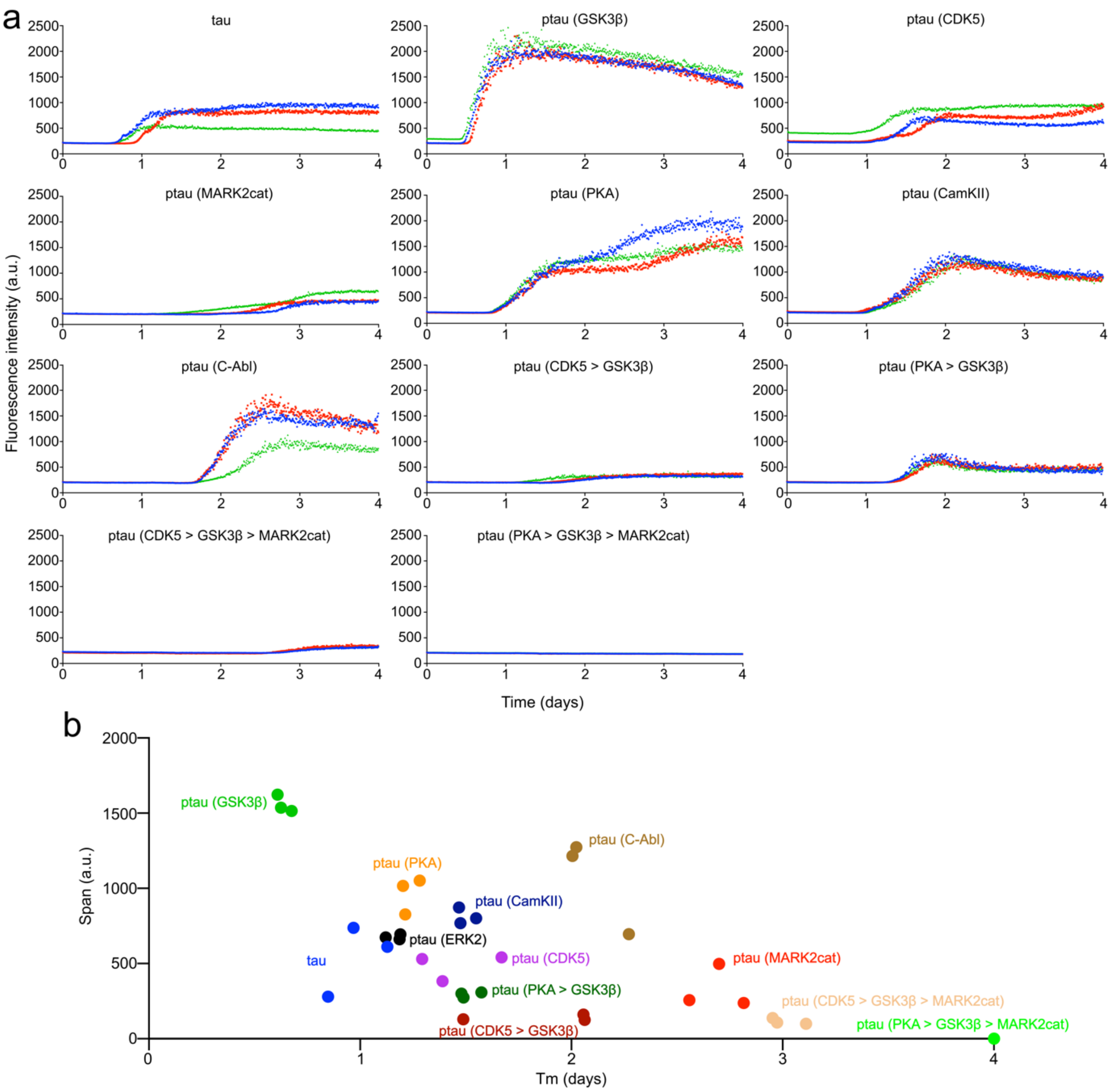


**Supplementary Fig. 3 | Aggregation kinetics of different phosphorylated tau samples. a,** Aggregation kinetics of three independent samples of 25 µM unmodified tau and tau phosphorylated by different kinases. **b,** ThT-intensity span vs. half time of aggregation (Tm) of unmodified and phosphorylated tau proteins.

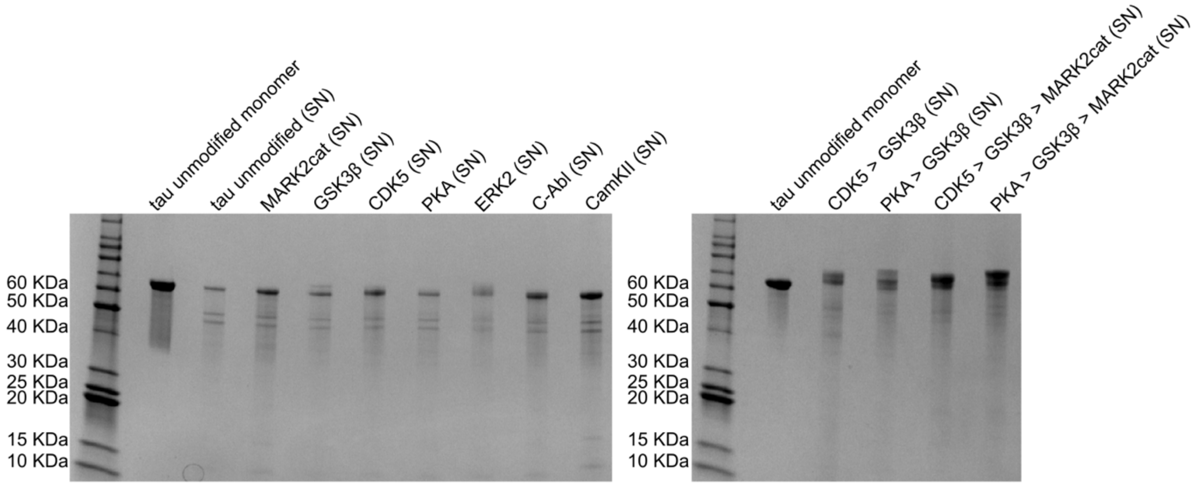


**Supplementary Fig. 4 | SDS-PAGE gel to determine the amount of aggregated protein.** SDS-PAGE gel of unmodified tau monomer and supernatant (SN) (after pelleting down the fibrils) of unmodified and different phosphorylated tau samples. The fibril samples were collected after four days of aggregation. The amount of aggregated protein was calculated by comparing the intensity of the supernatant (SN) band to the tau monomer band.

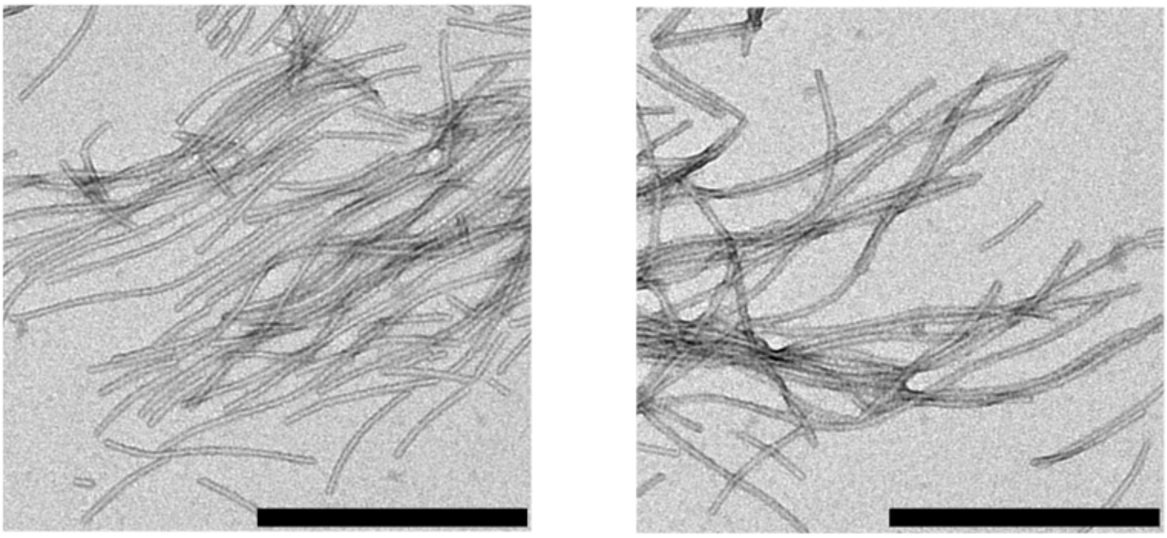

**Supplementary Fig. 5 │ Negative-stain EM images of unmodified tau fibril.** Scale bar, 500 nm.

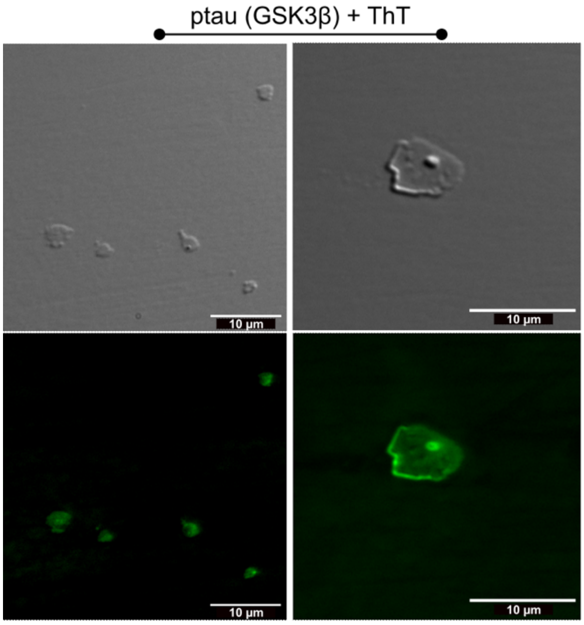


**Supplementary Fig. 6│ ThT staining of condensates formed by GSK3β-phosphorylated tau.** DIC (top panel) and fluorescence microscopy (bottom panel) of condensates formed by GSK3β-phosphorylated tau. Scale bars, 10 μm.

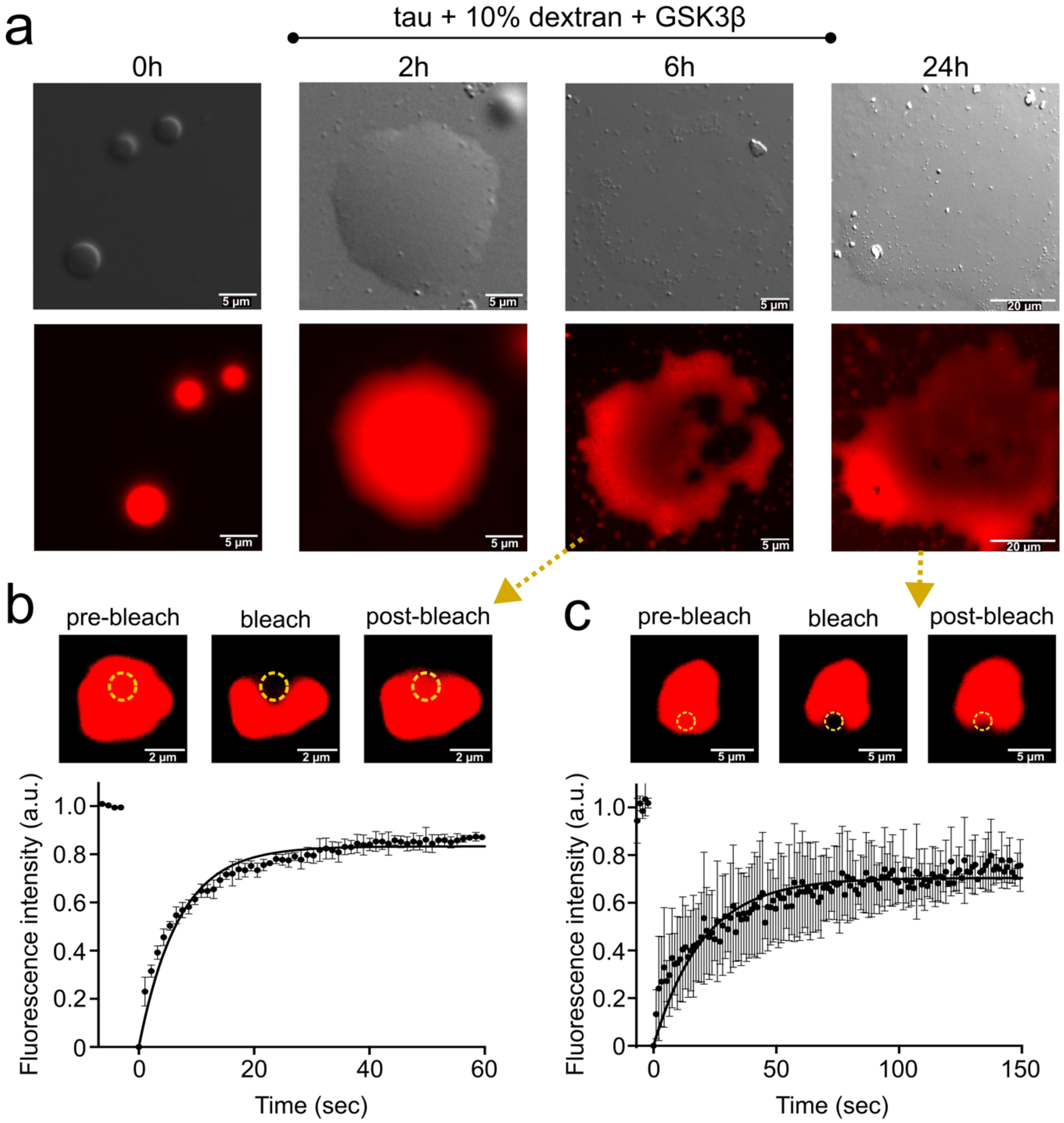


**Supplementary Fig. 7 | Incubation of tau droplets in the presence of GSK3β but without ATP. a,** DIC and fluorescence microscopy of tau droplets induced by the addition of 10 % dextran at room temperature in 25 mM HEPES, 5 mM $MgCl_2$, pH 7.2 buffer in the presence of 0.02 mg/ml unlabeled GSK3β (without ATP, i.e. under the conditions where no phosphorylation occurs). The sample was incubated for a duration of 24 hours. Micrographs are representative of three independent biological replicates. **b**, FRAP experiment of the tau condensates in the presence of GSK3β after incubation for six hours. Error bars represent the std of averaged three curves for each time point. Representative micrographs of the condensate before bleaching, after bleaching, and at the end of recovery are displayed (top panel). **c**, FRAP experiment of the tau condensates in the presence of GSK3β after incubation for twenty-four hours. Error bars represent the std of averaged three curves for each time point. Representative micrographs of the condensate before bleaching, after bleaching, and at the end of recovery are displayed (top panel).

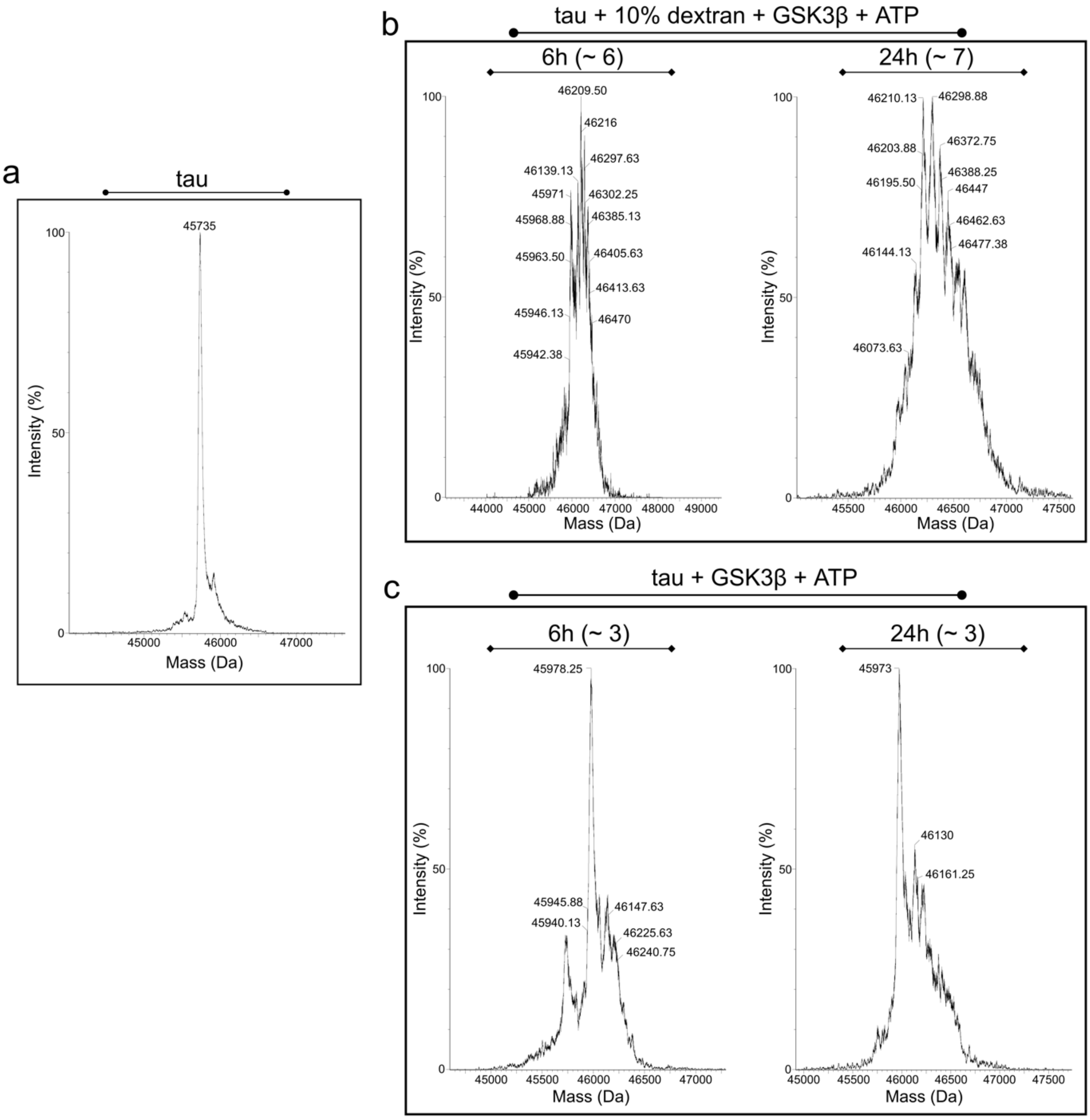


**Supplementary Fig. 8│ Mass spectrometry of tau phosphorylation. a,** Mass spectra of monomeric tau protein. **b**, Mass spectra of tau droplets induced by the addition of 10 % dextran at room temperature in 25 mM HEPES, 5 mM $MgCl_2$, pH 7.2 buffer in the presence of 0.02 mg/ml unlabeled GSK3β and 1 mM ATP after incubation for six hours (left) and twenty-four hours (right). The average degree of phosphorylation is indicated within the brackets. **c**, Mass spectra of tau monomer at room temperature in 25 mM HEPES, 5 mM $MgCl_2$, pH 7.2 buffer in the presence of 0.02 mg/ml unlabeled GSK3β and 1 mM ATP after incubation for six hours (left) and twenty-four hours (right). The average degree of phosphorylation is indicated within the brackets.

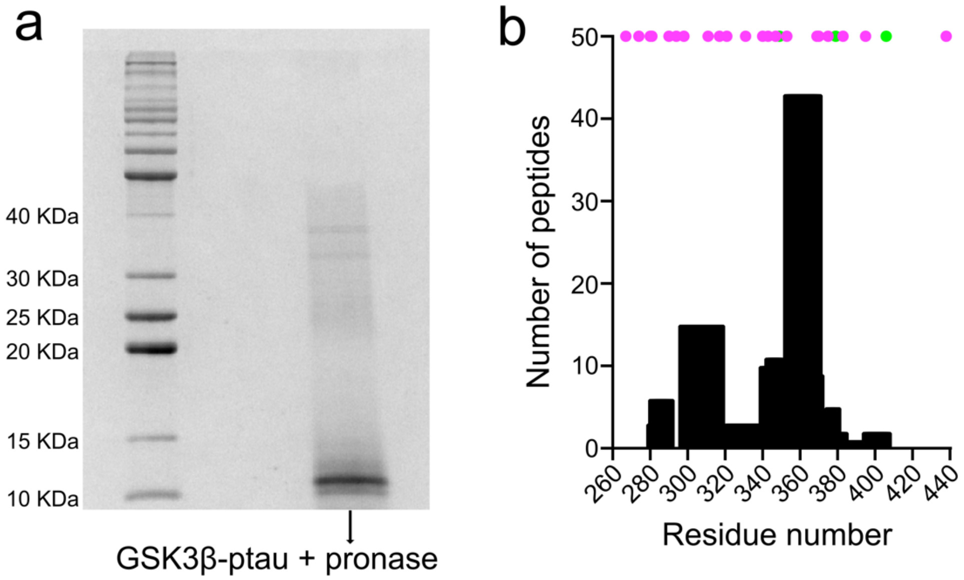


**Supplementary Fig. 9| Determination of the location of the rigid core of GSK3β-phosphorylated tau fibrils. a,** SDS-PAGE gel of pronase-digested GSK3β-phosphorylated tau fibrils. **b**, Numbers of peptides detected by mass spectrometry. The position of lysine and arginine residues are marked with purple and green dots, respectively.

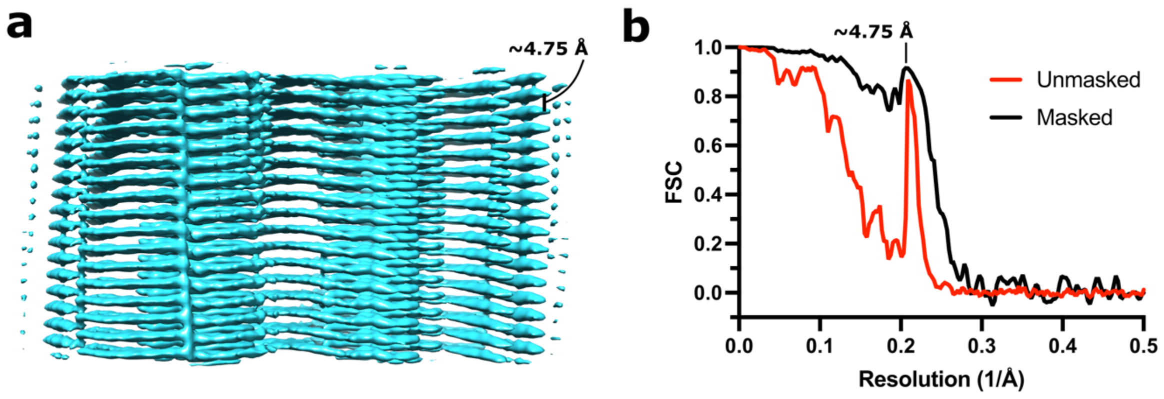


**Supplementary Fig. 10 | Cryo-EM of GSK3β-phosphorylated tau fibrils. a,** Cryo-EM density map of GSK3β phosphorylated tau fibrils from the side view, showing the high z-axis resolution. **b,** Fourier shell correlation (FSC) curves of GSK3β phosphorylated tau fibril maps. FSC curves between two independently refined masked (black) and unmasked (red) half-maps. The final resolution estimated from the value of the FSC curve for two independently refined masked half-maps at 0.143 is 3.85 Å. The peak at 4.75 Å from the high z-axis resolution results in an overestimation of the resolution.

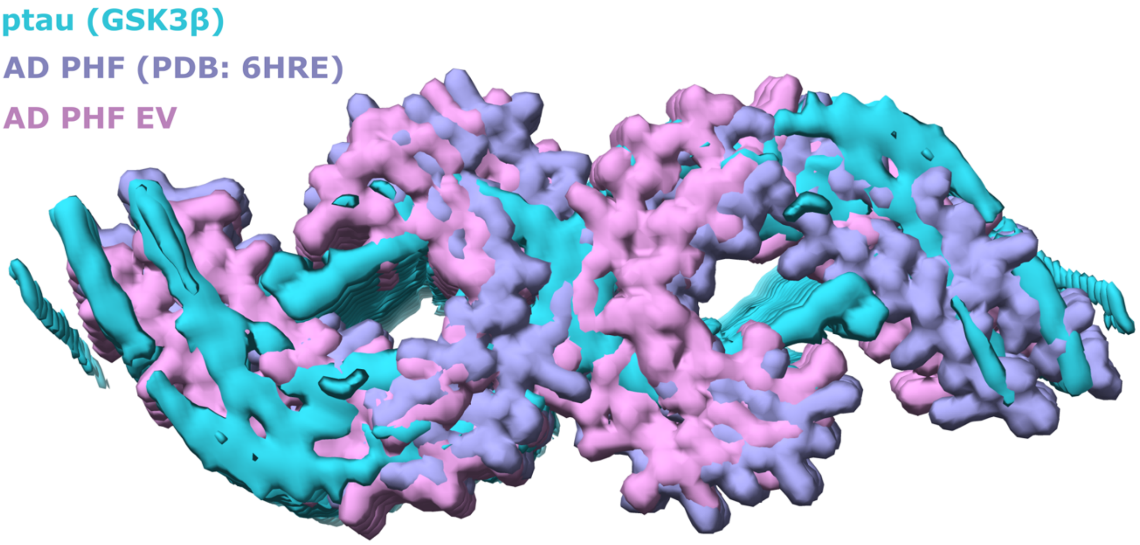


**Supplementary Fig. 11 │Superposition of the cryo-EM density maps of different tau fibrils.** Cryo-EM density map of the GSK3β-phosphorylated tau fibrils (cyan surface) compared with the PHFs from sporadic Alzheimer's disease brain (dark blue surface; PDB id 6HRE) and the PHFs extracted from extracellular vesicles (EV) of Alzheimer's disease brain (pink surface) (ref #48).

**Supplementary Table 1 | Cryo-EM collection and reconstruction data.**

| | |
|---|---|
| **Data collection** | |
| Microscope | Titan Krios G4 |
| Voltage (kV) | 300 |
| Detector | Falcon 4i |
| Pixel size (Å) | 0.934 |
| Defocus range (μm) | -0.9 to -1.9 |
| Exposure time (s) | 2.7 |
| Total dose ($e^-$/Å$^2$) | 40 |
| **Reconstruction** | |
| Picked segments | 1,315,404 |
| Box width (pixels) | 400 |
| Inter-box distance (pixels) | 18 |
| Final segments | 21,444 |
| Final resolution (Å)[a] | 3.85 (~5) |
| Sharpening B-factor (Å$^2$) | -26.43 |
| Symmetry imposed | C2 |
| Helical rise (Å) | 4.77 |
| Helical twist (°) | -0.58 |

[a] The resolution was estimated from the value of the FSC curve for two independently refined half-maps at 0.143. The resolution is overestimated because of the high resolution in the Z-axis. In brackets is shown the approximated real resolution.
